# Global warming readiness: Feasibility of enhanced biological phosphorus removal from wastewater at 35°C

**DOI:** 10.1101/2021.02.08.429585

**Authors:** Guanglei Qiu, Yingyu Law, Rogelio Zuniga-Montanez, Yang Lu, Samarpita Roy, Sara Swa Thi, Huiyi Hoon, Thi Quynh Ngoc Nguyen, Kaliyamoorthy Eganathan, Xianghui Liu, Per H. Nielsen, Rohan B.H. Williams, Stefan Wuertz

## Abstract

Recent research has shown enhanced biological phosphorus removal (EBPR) from municipal wastewater at warmer temperatures around 30°C to be stable in both laboratory-scale reactors and full-scale treatment plants. In the context of a changing climate, the feasibility of EBPR at even higher temperatures is of interest. We operated two lab-scale EBPR sequencing batch reactors with alternating anaerobic and aerobic phases for over 300 days at 30°C and 35°C, respectively, and followed the dynamics of the communities of phosphorus accumulating organisms (PAOs) and competing glycogen accumulating organisms (GAOs) using a combination of 16S rRNA gene metabarcoding, quantitative PCR and fluorescent in-situ hybridization analyses. Stable and nearly complete P removal was achieved at 30°C; similarly, long term P removal was stable at 35°C with effluent PO_4_^3−^-P concentrations < 0.5 mg/L on half of all monitored days. Diverse and abundant *Ca.* Accumulibacter amplicon sequence variants were closely related to those found in temperate environments, suggesting that EBPR at this temperature does not require a highly specialized PAO community. The slow-feeding strategy used effectively limited the carbon uptake rates of GAOs, allowing PAOs to outcompete GAOs at both temperatures. *Candidatus* Competibacter was the main GAO, along with cluster III *Defluviicoccus* members. These organisms withstood the slow-feeding regime, suggesting that their bioenergetic characteristics of carbon uptake differ from those of their tetrad-forming relatives. This specific lineage of GAOs warrants further study to establish how complete P removal can be maintained. Comparative cycle studies at two temperatures for each reactor revealed higher activity of *Ca*. Accumulibacter when the temperature was increased from 30°C to 35°C, suggesting that the stress was a result of the higher carbon (and/or P) metabolic rates of PAOs and GAOs, the resultant carbon deficiency, and additional community competition. An increase in the TOC to PO_4_^3-^-P ratio (from 25:1 to 40:1) effectively eased the carbon deficiency and benefited the proliferation of PAOs. In general, the slow-feeding strategy and sufficiently high carbon input benefited a high and stable EBPR at elevated temperature and represent basic conditions for full-scale applications.

## 1 INTRODUCTION

Enhanced biological phosphorus removal (EBPR) is a process widely employed for P removal in municipal wastewater treatment plants (WWTPs), not only because it is relatively cheap and sustainable, but because it is suitable for downstream nutrient recovery. Despite the advantages offered by the EBPR process, its application in warm climates has long been considered unfavorable, largely owing to observations that the functional bacteria (i.e., polyphosphate accumulating organisms, PAOs) in the process were outcompeted by their rivals (glycogen accumulating organisms, GAOs) at temperatures above 25°C (Whang and Park, 2002; Panswad et al., 2003; Lopez-Vazquez et al., 2009). Deterioration of EBPR performance and the dominance of GAOs were reported at fullscale WWTPs from regions with an average yearly temperature above 25°C (Wong et al., 2005; Barnard and Steichen, 2006; Cao, 2011). Seasonal deterioration in EBPR performance as wastewater temperature increased during the summer has also been reported (Gu et al. 2005). In short-term batch tests, the maximum acetate uptake rates of *Candidatus* Accumulibacter - PAOs and *Candidatus* Competibacter - GAOs were similar at temperatures < 20°C (within the tested temperature range of 5-30°C, Brdjanovic et al., 1997). With increasing temperature, the acetate uptake rates of *Ca.* Competibacter continued to increase following the extended Arrhenius equation (Lopez-Vazquez et al. 2007). However, the rates of *Ca.* Accumulibacter remained relatively constant in the range of 20 - 30°C. Additionally, significantly increased anaerobic maintenance coefficients (exceeding those of *Ca.* Competibacter at >30°C) were observed for *Ca.* Accumulibacter. These results helped explain the EBPR instability reported in WWTPs treating warm effluents (Lopez-Vazquez et al. 2007). However, these experiments were performed by exposing the enrichment culture obtained at 20°C to other temperature conditions, and hence it is not clear if the observed short-term effects were partially a stress response by *Ca.* Accumulibacter to acute temperature changes.

On the other hand, there have been efforts to explore the long-term feasibility and stability of EBPR at temperatures >25°C. Robust EBPR activity was obtained at 30°C for an extended period (>100 days) by applying a short EBPR cycle (Freitas et al. 2009). Selectively removing biomass from the top of the sludge bed allowed to maintain complete P removal for nearly 3 months in an aerobic granular sludge system operated at 30°C (Winkler et al. 2011). Recently, long-term EBPR (160 days) stability has been demonstrated in a lab-scale reactor at 28°C, without applying a short EBPR cycle or selective sludge removal (Ong et al., 2014). However, as the temperature increased to 32°C, EBPR activities were significantly compromised, with a reduction in the average abundance (based on qPCR analyses) of *Ca.* Accumulibacter and concurrent increase in the GAO population. Using acetate as a sole carbon source, the predominant PAO in the system was *Ca.* Accumulibacter clade IIF. It was hypothesized that this specific clade of *Ca.* Accumulibacter might have a better tolerance to high temperature. Shen et al. (2017) further showed that having multiple anaerobic/aerobic stages in one EBPR aided EBPR at 30°C. Acetate as a carbon source resulted in higher process stability compared to propionate.

In addition to these laboratory-scale studies, which included either special process operation control or the presence of a specifically enriched bacterial PAO community, there are reports of stable EBPR in full-scale systems in warm climates (Sayi-Ucar et al., 2015; Law et al. 2016; Cao et al., 2017). The presence of both GAOs and PAOs had no apparent effect on long-term stability of EBPR at 30°C (Law et al., 2016). A recent field study in Singapore further showed a wide range of PAOs in plants operating at water temperatures from 28-32°C exhibiting high micro-diversity in carbon usage, which might be important for the observed EBPR stability in these full-scale plants (Qiu et al., 2019).

Even higher water temperatures are expected due to global warming. The global surface temperatures are likely to rise up to 4.8°C by the end of this century in a highest emissions scenario (IPCC, 2018). More pronounced temperature increases have been observed in highly urbanized areas/regions; e.g., according to the Meteorological Service Singapore, temperatures in Singapore have risen at a rate (0.25°C in each decade) more than double that of the global average during the period of 1948 to 2015 (Meteorological Service Singapore, 2021).

The aim of this work was to explore the feasibility of EBPR at 35°C. Two lab-scale sequencing batch reactors (SBRs) were operated in parallel at 30°C and 35°C, respectively, for over 300 days. A slow-feeding strategy was employed with a mixture of acetate and propionate (a molar ratio of 8.4:1) as carbon source to mimic the operation and carbon sources conditions typically found in the field (Qiu et al., 2019). Consecutive cycle studies involved switching temperatures between 30°C and 35°C to investigate the effect of temperature on carbon and P transformation kinetics and stoichiometry. The fine-scale dynamic of the bacterial community was monitored using fluorescent in-situ hybridization (FISH), 16S rRNA gene metabarcoding and quantitative polymerase chain reaction (qPCR).

## 2 MATERIALS AND METHODS

### 2.1 Sequential batch reactor (SBR) operation

Two sequencing batch reactors (SBR) with a working volume of 1.59 L were inoculated with activated sludge from a local WWTP in Singapore (Qiu et al., 2019). A slow feeding strategy mimicking the actual feeding conditions in the full-scale plant was applied for reactor operation. Slow feeding has been shown to benefit the proliferation of PAOs at 22°C (Tu and Schuler, 2014). The strategy was showed to be effective in obtaining highly enriched *Ca.* Accumulibacter culture at high temperature (30°C, Qiu et al., 2020). Additionally, a mixture of acetate and propionate (with a molar ratio of around 8.4:1) representing the typical VFAs composition in full-scale plants (Qiu et al., 2019) was used as a carbon source. The SBRs were operated with 6-h cycles, including a 60-min feed, a 20-min anaerobic, a 180-min aerobic, and a 100-min settling/decant stage. In each cycle, 0.740 L of synthetic wastewater composed of 0.050 L of solution A (containing 1.02 g/L NH_4_Cl, 1.2 g/L MgSO_4_·7H_2_O, 0.01g/L peptone, 0.01g/L yeast extract, 3.92 g/L sodium acetate and 0.255 g/L propionic acid) and 0.69 L of solution B (34.6 mg/L K_2_HPO_4_·3H_2_O, 20.6 mg/L KH_2_PO_4_, 0.75 mg/L FeCl_3_·6H_2_O, 0.015 mg/L CuSO_4_·5H_2_O, 0.03 mg/L MnCl_2_, 0.06 mg/L ZnSO_4_, 0.075 mg/L CoCl_2_, 0.075 mg/L H_3_BO_3_, 0.09 mg/L KI and 0.06 mg/L Na_2_MoO_4_·2H_2_O) were introduced into the reactor, with a resultant TOC/P molar ratio of around 25:1. Two reactors were operated in parallel at 30°C (R30) and 35°C (R35), respectively. The HRT and SRT were 12.9 h and 25 d in both reactors. The pH was controlled at 6.80 - 7.50, with DO levels maintained at 0.8-1.2 mg/L during the aerobic phase. During Day 175-220, the sodium acetate and propionic acid concentration in solution A for R35 were increased by 1.6 time (with the corresponding TOC/P molar ratio increased to 40:1), which was further reduced to 1.2 times of their original values (TOC/P molar ratio= 30:1) from Day 216 onwards until the end of the experiment.

### 2.2 Consecutive cycle study

A consecutive cycle study was performed on Days 55, 77, 105, 203 and 294 to investigate the effect of temperature on the carbon (C) and P cycling kinetics and stoichiometry of the EBPR community. Cycle studies were done in adjacent cycles in both reactor with the first SBR cycle operated at their respective original temperature, i.e., R30 at 30°C and R35 at 35°C. In the following cycle, the temperature of the two reactors was switched, i.e., R30 at 35°C, and R35 at 30°C. This setting allows the test of the temperature effects in both reactors but effectively deduct any bias potentially arising from the acute response of the community to temperature changes. Filtered (through 0.45μm membrane filters) and activated sludge samples were collected at regular time intervals for PO_4_^3-^-P, VFAs, polyhydroxyalkanoate (PHA) and glycogen analyses.

### 2.3 Chemical analysis

PO_4_^3-^-P were measured using test kits (HACH, CO, USA) following the Standard Methods (APHA, 1999). Mixed liquor suspended solid (MLSS) and mixed liquor suspended volatile solid (MLVSS) concentrations were determined according to the Standard Methods (APHA, 1999). Acetate and propionate concentrations were measured using a gas chromatograph (Prominence, Shimadzu, Japan) equipped with a DB-FFAP column (30×0.25 mm) (Agilent Technology, U.S.) and a flame ionization detector. PHA analyses were performed using a gas chromatograph (Prominence, Shimadzu, Japan) equipped with a DB-5MS Ultra Inert column (30×0.25 mm) (Agilent Technology, CA, USA) and an FID detector (Oehmen et al. 2005a). Glycogen analyses were carried out by measuring glucose after acid digestion of the freeze-dried sludge as described by Kristiansen et al. (2013).

### 2.4 Fluorescence in situ hybridization (FISH)

Activated sludge samples collected from both reactors during the operation was immediately fixed using paraformaldehyde (PFA, to a final concentration of 4%) at 4°C for 2 h. The fixed samples were washed with 1×phosphate-buffered saline (PBS) solution and stored in a mixture of 1×PBS and ethanol (1:1) at −20°C before FISH analysis. Microorganisms of interest were detected using EUBmix (EUB338, EUB338II and EUB338III) probe targeting most bacteria (Daims et al., 1999), PAOmix (PAO651, PAO462 and PAO846) targeting *Ca.* Accumulibacter (Crocetti et al., 2000), GAOmix (GAO431 and GAO989) targeting *Ca.* Competibacter (Crocetti et al., 2002; Kong et al., 2002), and TFOmix (DF218 and DF618) (Wong et al., 2004) and DFmix (DF988, DF1020) (Meyer et al., 2006) targeting cluster I and cluster II *Defluviicoccus*, respectively (DF988 was also showed to target *Nostocoida limicola-like* cluster III members, Nittami et al., 2009). FISH images were collected using a LSM780 confocal laser scanning microscope (CLSM) equipped with a laser light source (Carl Zeiss, German). The Zen (black edition, Carl Zeiss, German) software was used for image processing and output.

### 2.5 DNA extraction, 16S rRNA gene metabarcoding and qPCR

Genomic DNA was extracted using the Fast SPIN Kit for Soil samples (MP Biomedicals, CA, USA) (Albertsen et al. 2015). Bacterial 16S rRNA gene metabarcoding was carried out, targeting the V1-V3 region with primer set: 27F (5’-AGAGTTTGATCCTGGCTCAG-3’) and 534R (5’-ATTACCGCGGCTGCTGG-3’). Extracted genomic DNA were subjected to 16S rRNA gene metabarcoding analysis by Miseq (Illumina, CA, USA) at the Australian Centre for Ecogenomics (Bribane, Australia). The generated data was analyzed using the DADA2 pipeline (Version 1.12, Callahan et al., 2016; 2017) using MiDAS 3.6 (McIlroy et al. 2015; Nierychlo et al., 2020) as reference for taxonomy assignment.

Additionally, quantitative PCR (qPCR) was used to analyze the clade level distribution of *Ca.* Accumulibacter, according to He et al. (2007).

### 2.6 Statistical analysis

Statistical analyses were performed using R version 3.6.3 (R Core Team, 2020). The alpha diversity calculation and PCoA analysis were performed using R packages “hillR” version 0.5.0 (Chao et al., 2014) and “vegan” version 2.5-6 (Oksanen et al., 2019). Microbial ecological network analysis was performed using Cytoscape version 3.7.2 with the app package CoNet version 1.1.1 (Shannon et al., 2003)

## 3 RESULTS AND DISCUSSION

### 3.1 EBPR is feasible at 35°C

Two laboratory-scale SBRs were operated in parallel to investigate the feasibility of EBPR at 35°C. A slow feeding strategy was employed to maintain low in-tank acetate concentrations mimicking the conditions typically found in the field. Low acetate concentration has been shown to selectively benefit *Ca.* Accumulibacter over *Defluviicoccus-GAOs* (Tu and Schuler, 2013), owing to the higher ability of *Ca.* Accumulibacter to scavenge low-concentrations of acetate via an acetate-proton symporter (ActP) driven by the proton motive force (PMF) (Saunders et al., 2007; Burow et al., 2008; Qiu et al., 2020). We hypothesized that the slow-feeding strategy further benefits EBPR at high temperature by effectively limiting the carbon uptake rates of *Ca.* Competibacter. Additionally, a mixture of acetate and propionate was used as carbon source. Mixed carbon sources were shown to selectively benefit *Ca.* Accumulibacter over GAOs (*Ca.* Competibacter and *Defluviicoccus*) in a modeling work (Lopez-Vazquez et al., 2009), since *Ca.* Competibacter and *Defluviicoccus* were inefficient in using propionate and acetate, respectively (Oehmen et al., 2005b; Oehmen et al., 2006), but *Ca.* Accumulibacter could take up both carbon sources at similarly high rates (Oehmen et al., 2005b).

With influent TOC and PO_4_^3-^-P concentrations of 86 mg/L (COD, 230 mg/L) and 8.8 mg/L, respectively, (corresponding to a TOC/P molar ratio of 25:1), R30 (the reactor operated at 30°C) showed high and stable EBPR performance over 300 days (Fig. 1A). P-release values were maintained at >35 mg/L. Complete P removal was achieved on most days. A technical failure occurred on Day 35, when no carbon source was supplied for 2 days. Good EBPR activity was restored immediately after the system operation had returned to normal, showing the high resilience of the system to unexpected disturbances. Previous studies reported that EBPR activity at temperatures up to 32°C for up to 80 days, but with largely suppressed EBPR activities (Ong et al., 2014). Similarly, Shen et al. (2017) observed significant fluctuating EBPR performance although complete P removal was achievable in periods at 30°C, using a multi-cycle operation strategy. The combined strategy of slow feeding and mixed carbon sources (acetate/propionate molar ratio of 8.4:1) used in this work likely benefited a high and stable EBPR performance at 30°C. These results also explain why EBPR at 30°C is readily achieved in the field without specific process control (Law et al. 2016; Cokro et al. 2019) compared to laboratory-scale studies.

**Figure 1.**
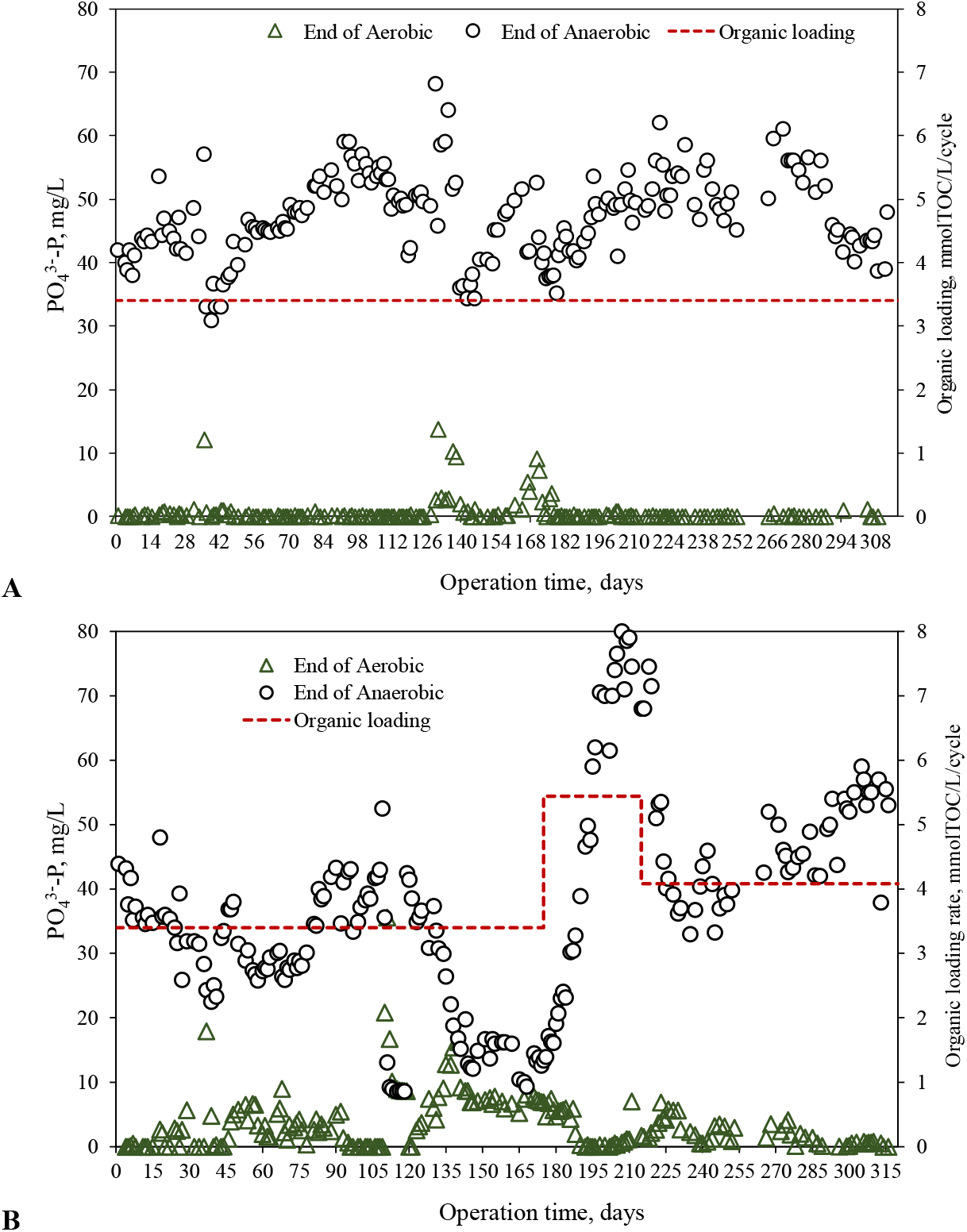
Phosphorus removal performance in the EBPR reactors (**A**. R30) at 30°C and (**B**. R35) 35°C. Slow-feeding regime was used with the influent wastewater being introduced into reactors at fix speeds in 60 min. The carbon source was a mixture of acetate and propionate (with a molar ratio of around 8.4:1). The resultant TOC/P molar ratio was 25:1. The TOC/P molar ratio in R35 was increased to 40:1 during Day 175-220, which was further reduced to 30:1 from Day 220 onwards.

Complete P removal was also achieved in R35 (the reactor operated at 35°C) in the first 2 weeks of operation (Fig. 1B). After that, incomplete P removal occurred occasionally, but appreciable P-release and -uptake activities (>25 mg/L) were still maintained (with P removal efficiencies >70%). The same technical fault that affected R30 occurred on Day 35, and the following 2-day starvation period had no lasting impact on EBPR performance because complete P removal was restored within 2 days. EBPR was maintained during Days 91-109, until there was a sudden drop in the P-release and -uptake activities from around 40 to 0 mg/L within a day. The reason for this decrease in EBPR activity is unknown but may have been related to phage predation. Phage-related genes have been identified in *Ca.* Accumulibacter genomes (Garcia Martin et al., 2006; Flowers et al., 2013) and EBPR failure caused by bacteriophages has been reported (Barr et al., 2010; Motlagh et al., 2015). The reactor (R35) did not recover on its own from this complete loss of EBPR activity (Days 109-119, Fig. 1B). Reseeding with waste sludge collected from R30 (half the amount of biomass in R35 was removed and replaced with waste sludge from R30) allowed a restoration of EBPR activities, although at a lower level (P-release and -uptake values around 10 mg/L, Days 135-175). An increase in the influent TOC concentration to 137 mg/L (1.6 times the original TOC, with a corresponding increase in the TOC/P ratio to 40:1) led to a rapid increase in EBPR activities. Complete P removal was achieved on Day 187 and maintained until Day 219 when the influent TOC was reduced to 103 mg/L (1.2 times the original TOC and a corresponding TOC/P ratio of 30:1). After a period of low-level fluctuation in the EBPR performance, near-complete P removal was achieved again on Day 285, which was maintained until the end of the experiment (Day 316). In general, although the overall performance and system stability were lower compared to R30, appreciable EBPR activities were maintained in R35 over an extended time period (with an average effluent PO_4_^3-^-P of 1.96 mg/L during the entire operation excluding the period of unexpected collapse and subsequent recovery). Near complete P removal was achieved (with effluent PO_4_^3-^-P < 0.5 mg/L) more than half the time. The reactor performance recovered quickly from a disturbance caused by a technical fault resulting in starvation and was also rescued after an unexplained system failure by inoculation with sludge from R30, suggesting that 35°C is not an insurmountable obstacle for EBPR. Previous work had demonstrated the feasibility of EBPR at 32°C (Freitas et al., 2009; Ong et al., 2014; Shen et al., 2017), and Panswad et al. (2003) showed that EBPR activities were detectable at 32.5°C but ceased at 35°C. Hence the present study is the first report of sustained EBPR at this temperature.

### 3.2 Community composition and dynamics

To understand the relationship between PAO and GAO dynamics and EBPR performance at high temperature the bacterial communities in both reactors were characterized via 16S rRNA gene metabarcoding, FISH and qPCR. An overview of the top 30 most abundant genus during the longterm operation of two reactors are showed in the Supplementary Material, Fig. S1.

#### 3.2.1 Polyphosphate accumulating organisms in the reactors are closely related to these commonly found in temperate systems

Thirty-nine amplicon sequence variants (ASVs, each ASV was uniquely numbered, the same ASV number indicates the same sequence) were affiliated with the *Ca.* Accumulibacter lineage, 27 and 28 ASVs of which occurred in R30 and R35, respectively (Fig. 2A and B). The two reactors shared 16 ASVs, with 9 of them (ASVs 11, 23, 28, 32, 73, 36, 38, 87, 353) being detected in >4 samples in both reactors. These 9 ASVs also represented the dominant *Ca.* Accumulibacter species during the operation of the two reactors. Among these 9 core ASVs, 6 (ASVs 23, 28, 32, 73, 38, 353) were originally present in the seed sludge (there were 7 ASVs in total in the seed sludge, with relative abundance values ranging from 0.06 to 0.28%). The remaining 3 ASVs (11, 73 and 87) started to emerge in both reactors from Day 5, implying that they were probably also present in the seed sludge but at relative abundances under the detection limit. The relative abundance values of all ASVs in the seed sludge increased in both reactors at the beginning of the operation, except for ASV 1033 (the 7th ASV, with a relative abundance value of 0.07%) which decreased and disappeared in both reactors within 30 days. These results suggested that the operation strategy employed in this study effectively retained the majority of the *Ca.* Accumulibacter species originally present in the full-scale sludge, which allowed them to proliferate and predominate at different stages during the reactor operation. The diverse *Ca.* Accumulibacter community enriched in the two reactors is also considered a key to the stable and reliable EBPR activities observed in the long term. Additionally, these results demonstrated that most of the *Ca.* Accumulibacter species in the full-scale sludge could effectively survive an elevated temperature of 35°C.

**Figure 2.**
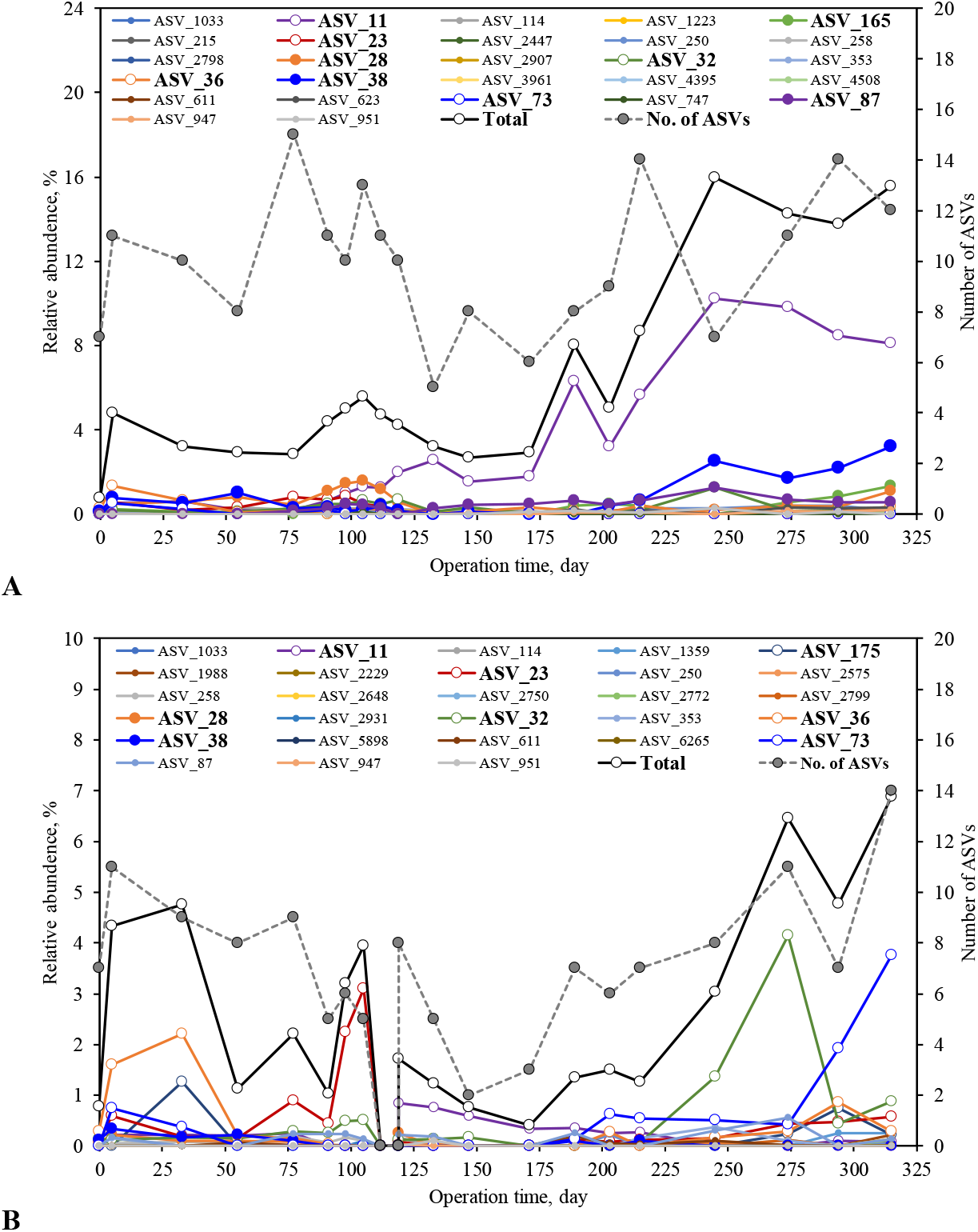
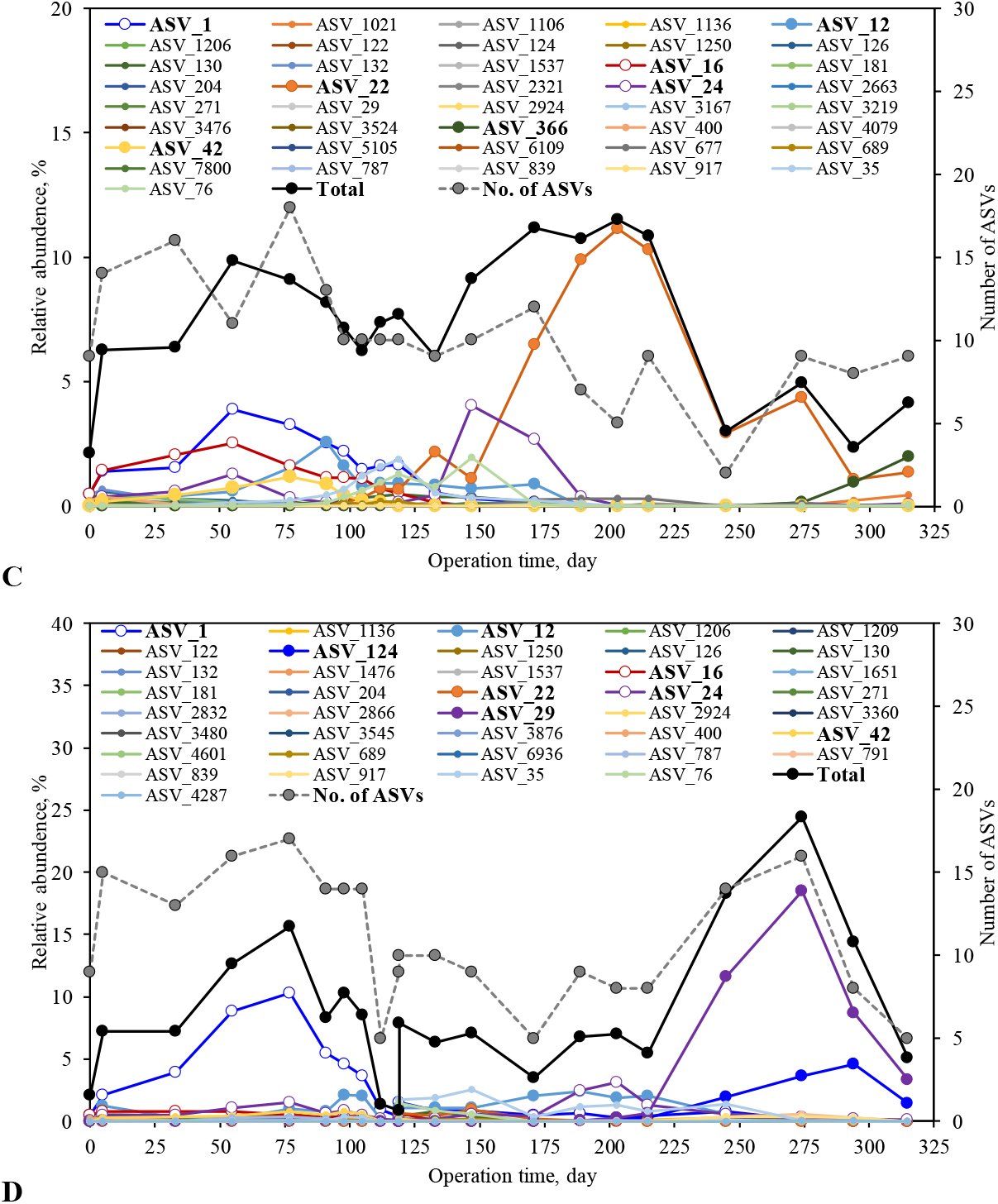
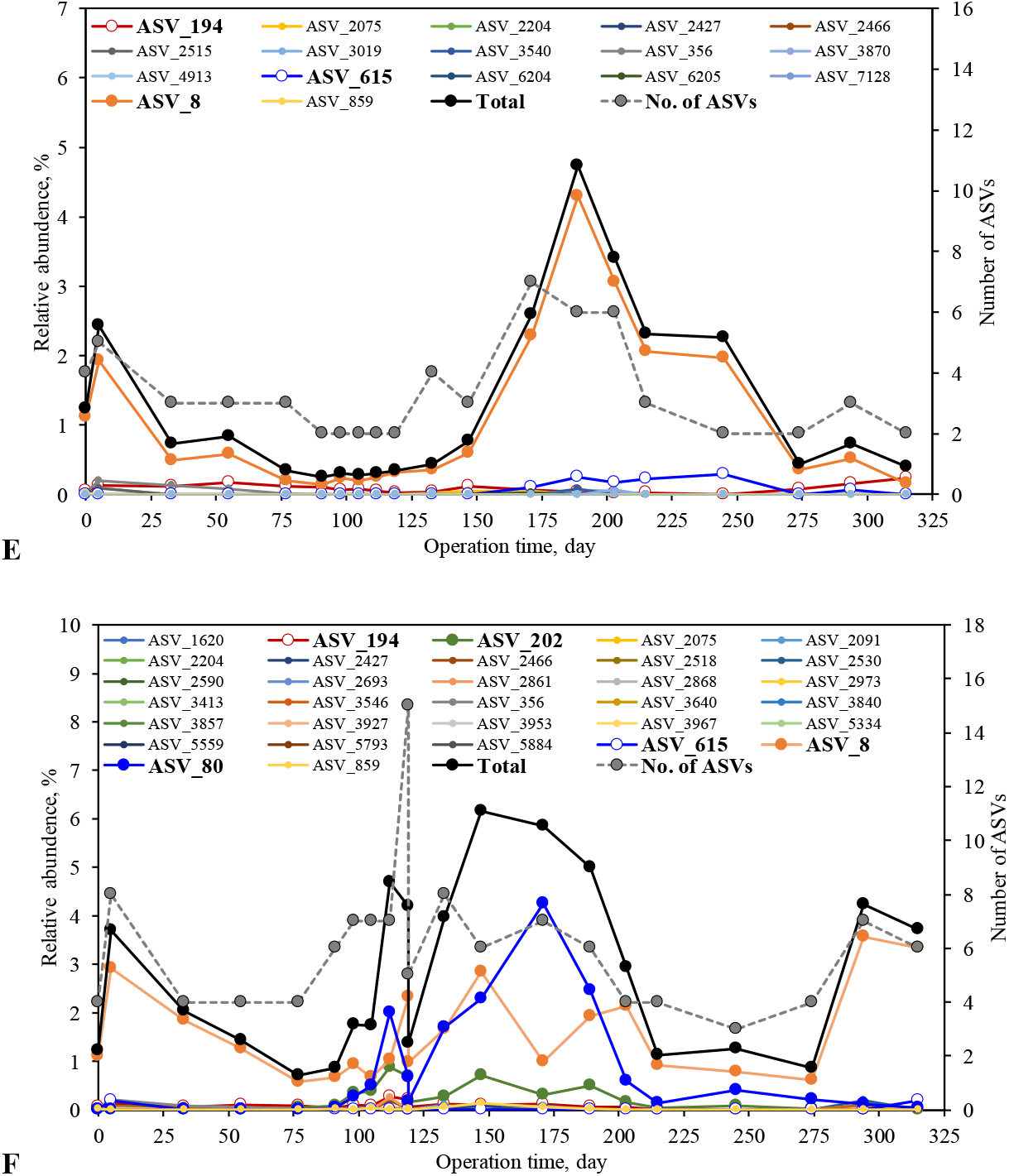
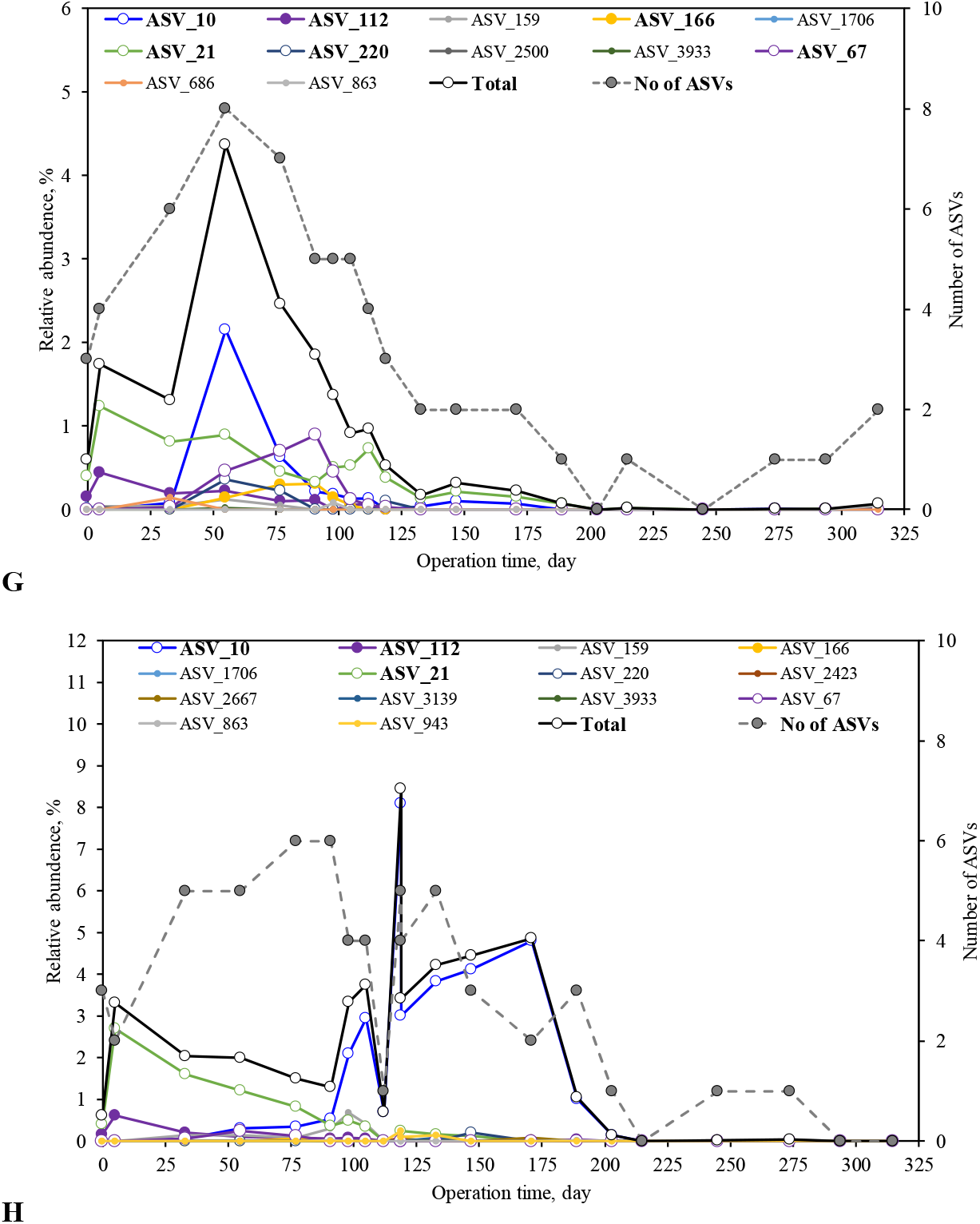
Community composition and dynamics revealed by 16S rRNA gene amplicon sequencing: (**A**) *Ca.* Accumulibacter in R30; (**B**) *Ca.* Accumulibacter in R35; (**C**) *Ca.* Competibacter in R30; (**D**) Ca. Competibacter in R35; (**E**) *Defluviicoccus* in R30; (**F**) *Defluviicoccus* in R35; (**G**) *Dechloromonas* in R30; (**H**) *Dechloromonas* in R35.

Despite the extended presence of the core *Ca.* Accumulibacter ASVs, their relative abundance values and status of predominance were largely different in the two reactors, with significant community dynamics within each reactor during the operation (Fig.2A, 2B and Supplementary material Fig.S1). In R30, ASVs 11, 23, 28, 32, 36, 38 were present at comparable relative abundances ranging from 0.07% to 1.57%. Later, ASV 11 started to dominate in R30 until the end of operation. ASV 11 is closely related (100% identity) to *Ca.* Accumulibacter clade IIF strain SCELSE-1 (Qiu et al. 2020). Similarly, Ong et al. (2014) reported that the dominating *Ca.* Accumulibacter clade in a bioreactor at 32°C belonged to clade IIF. It seems that certain metabolic characteristics enable members of the clade to dominate the high temperature at around 30°C. *Ca.* For example, Accumulibacter clade IIF strain SCELSE-1 can use a wide range of amino acids for EBPR (Qiu et al., 2020). Genomic analysis also suggested that clade IIF member genomes encode enzymes necessary for the conversion of ethanol to acetate (Skennerton et al., 2014). Their versatility in carbon usage and hence the potential to benefit from various metabolites from other community members could explain their predominance in the bioreactors studied here. Notably, ASV 11 was less abundant in R35 than in R30 throughout the operation. The sudden disruption of EBPR activities in R35 and subsequent inoculation with R30 sludge on Day 119 presented an opportunity to test the growth potential of ASV11 at 35°C. Although this ASV was not completely eliminated from the reactor, its relative abundance decreased to <0.1% over time, implying that ASV 11 does not have a significant growth advantage over other ASVs at 35°C. Compared to R30, R35 exhibited dominance of a single ASV during each period of operation. ASV 36 prevailed during the first 50 days, followed by ASV 23 and ASV 11 and then ASV32 and ASV73. A whole-genus phylogenetic analysis suggested that these ASVs are widely distributed in the *Ca.* Accumulibacter lineage (Supplementary Material Fig. S2). ASV 36 is closely related (100% identity) to *Ca.* Accumulibacter species EF565158/EF565159 (sequences recovered from a full-scale WWTP at Durham in the U.S., He et al., 2007), which are clade IID members. ASV 32 is closely related (100% identity) to AB736230 (a sequence recovered from a lab-scale EBPR system in Japan, Satoh et al., 2013), which branched in the same lineage of EF565149 (from a full-scale WWTP in the U.S., He et al., 2007) and AF204248 (from a lab-scale SBR operated at 22°C in California, Crocetti et al., 2000), indicating that ASV 32 is probably a clade IIB member. ASV11 is closely related to *Ca.* Accumulibacter clade IIF stain SCELSE-1 (Qiu et al., 2020). ASV23 showed 100% identity to HQ506555 (from a full-scale WWTP plant in Korea, Kim et al., 2013a); and ASV73 is closely related to HQ158615 (form a lab-scale SBR reactor, Zhu and Chen, 2011) and AB736228 (Satoh et al., 2013). The clade identities of these two ASVs are not clear. Additionally, the dominant ASV (87) in R30 is closely affiliated (100% identity) with a clade IIF member (JQ 726366) in an SBR reactor operated at 20°C in Korea (Kim et al., 2013b). ASV28 is closely related to HQ010772 (100% identity, Ji and Chen, 2010), KJ807999 (100% identity, from a full-scale WWTP in China, Wang et al., 2014) and EF565160 (99.27% identity, He et al., 2007); and ASV38 is closely relate to JQ726361 (Kim et al., 2013b). Both ASVs are probably clade IIC members. qPCR analysis was employed to analyze the clade-level distribution and dynamics of *Ca.* Accumulibacter (Supplementary material Fig. S3). No clade I member was detected in either of the two reactors. The dynamics in the relative abundances of clades IIB, IIC and IID generally agree with the 16S rRNA gene-based identification of these dominant ASVs. These results suggested that, a diverse range of clade members were able to survive at 35°C. These dominated ASVs are not unique but also found in world-wide distributed laboratory- and full-scale EBPR systems in temperate climates, implying that successful EBPR at elevated temperatures does not require a highly specialized *Ca.* Accumulibacter community.

#### 3.2.2 Glycogen accumulating organisms dominated the reactors are not commonly detected in fullscale systems

Fifty-four sequences were affiliated with *Ca.* Competibacter, and 40 of them were found in each reactor (Fig. 2C and 2D). The overall trends were similar to those of *Ca.* Accumulibacter in the two reactors. Multiple ASVs (1, 12, 16, 24, 42) dominated R30 for the first 150 days. ASV 22 proliferated during the remainder of reactor operation until ASV 366 appeared towards the end of the experiment. Single ASVs dominated in R35 during each period of operation (ASV 1 for the first 100 days and ASV 29 and ASV 124 afterwards). Different from the core *Ca.* Accumulibacter ASVs, which were all found in the seed sludge, both ASV 29 and ASV 124 in R35 and ASV 22, which dominated RS30 at the later stage (from day 150 onward), were absent or undetected in the seed-sludge. They appeared only after Day 77. No closely related relatives of these ASVs were found in the NCBI database. The best matches were LR650365 (from Aalborg, Denmark, 94.12% identity to ASV 29), HQ467835 (from a full-scale WWTP in Korea, 96.73% identity to ASV 124), and DQ201883 (from a Danish WWTP, 96.73% identity to ASV 22). Their finer scale identity and metabolic characteristics need further characterization.

Together these results suggest that the *Ca.* Competibacter species that appeared in the two laboratory-scale reactors do not play a significant role in the field, supporting the previous hypothesis that laboratory-scale studies tend to overestimate the impact of GAOs on the EBPR process (Nielson et al., 2019). Despite the presence of diverse *Ca.* Competibacter communities in both reactors with high relative abundances, there was no significant deterioration of EBPR activity. Their increased total relative abundances were generally associated with lower P-release values (Fig.1, and Fig.2C and 2D), showing that they did compete with *Ca.* Accumulibacter for carbon uptake.

*Defluviicoccus*-related GAOs were detected in both systems, although at low relative abundance compared to *Ca.* Competibacter (Fig 2E and 2F). A total of 41 ASVs were detected in the two reactors, of which 18 occurred in R30 and 30 in R35. ASV 8 dominated in R30, with other ASVs staying at low levels throughout the operation. The dominant ASV in R35 was also ASV 8 before Day 112 (when EBPR collapsed in the system). Afterwards ASV80 and ASV202 increased and their relative abundances declined subsequently as the EBPR activity recovered to a high level. ASV 8 took over again in the last 100 days. ASV 8 is phylogenetically closely related (100% identity) to AB445107 (Nittami et al., 2009), which was found as a Cluster III member having a *“Nostocoida Limicola-like”* filamentous morphology. In-situ ecophysiology analysis revealed a GAO phenotype (McIlroy, et al., 2010). They were also widely found in full-scale EBPR systems (McIlroy, et al., 2010; Stokholm-Bjerregaard et al., 2017). FISH analysis confirmed their presence and morphology in the two reactors (Fig. 4). This ASV was also the dominant *Defluviicoccus* species (with a relative abundance value of 1.11% out of 1.23%) in the seed sludge. Our previous field work showed their high relative abundance in full-scale plants in Singapore (Qiu et al., 2019). ASV 80 showed 99.63% similarity to DQ250533, a cluster II member having the typical tetrad-forming morphology (Wong and Liu, 2007). FISH analysis allowed their detection in R35 together with ASV 202 (which are cocci in clumps) (Fig. 4J). ASVs 80 and 202 were either not present in the seed sludge or were below the detection level. Their proliferation in R35 was probably a result of the deteriorated EBPR activity and the simultaneous reduction in the relative abundance of *Ca.* Competibacter (Fig. 2D), which could have opened up an ecological niche. Tetrad-forming *Defluviicoccus* members were believed to be unable to compete with *Ca.* Accumulibacter under slow-feeding conditions (Tu and Schuler, 2013) due to their inability to effectively take up acetate at low concentrations (Burow et al., 2008). Their decrease after the restoration of the EBPR activities verifies this hypothesis. However, the ability of Cluster III member ASV 8 to co-exist with *Ca.* Accumulibacter and *Ca.* Competibacter in the long term under the slow-feeding condition suggested that they are probably metabolically different from their Cluster II relatives in terms of carbon uptake mechanisms. They seem significant to EBPR in full-scale systems. Apart from these well-characterized PAOs and GAOs, *Dechloromonas-related* sequences were also detected in both reactors (Fig. 3G and 3H). Sixteen AVSs occurred in the two systems with 10 of them in R30 and 12 in R35. ASVs in R30 showed high inter-lineage dynamics in the first 125 days. Alternating predominance was observed for ASVs 10, 67, 112 and 166. In R35, trends were similar to those for the *Ca.* Accumulibacter and *Ca.* Competibacter community, where a significant predominance of single ASVs (10 and 21) was observed during different stages of operation. Previous research showed that there are probably both PAO and GAO phenotypes within the *Dechloromonas* lineage (Kong et al., 2007; Ahn et al., 2007; Günther et al., 2009). Recent research also allowed the identification of two novel *Dechloromonas*-related PAOs in full-scale WWTPs (Petriglieri et al., 2020). In R35, ASVs 10 and 21 were present during days 108-119, when there was no detectable EBPR activity, suggesting that both ASVs acted like GAOs. We did not find well-characterized relatives of these ASVs in the database (the highest similarity score obtain was 97.45%, which is between ASV67 and *Candidatus Dechloromonas phosphatis* midas_s_96, Petriglieri et al., 2020), and the existing FISH probes did not detect these dominant ASVs (McIlroy et al. 2016). Their ecological roles in the systems need further confirmation.

**Figure 4.**
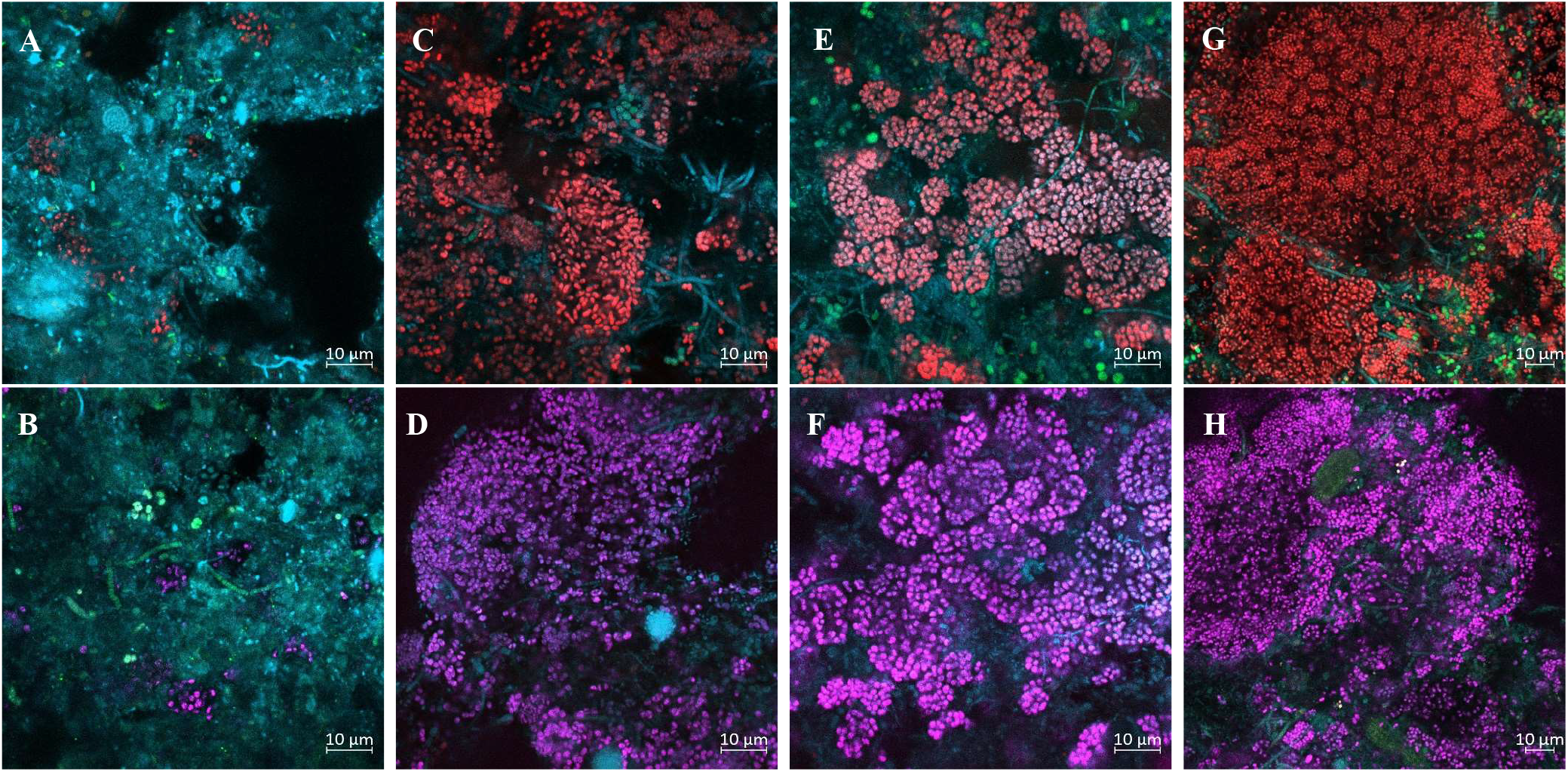
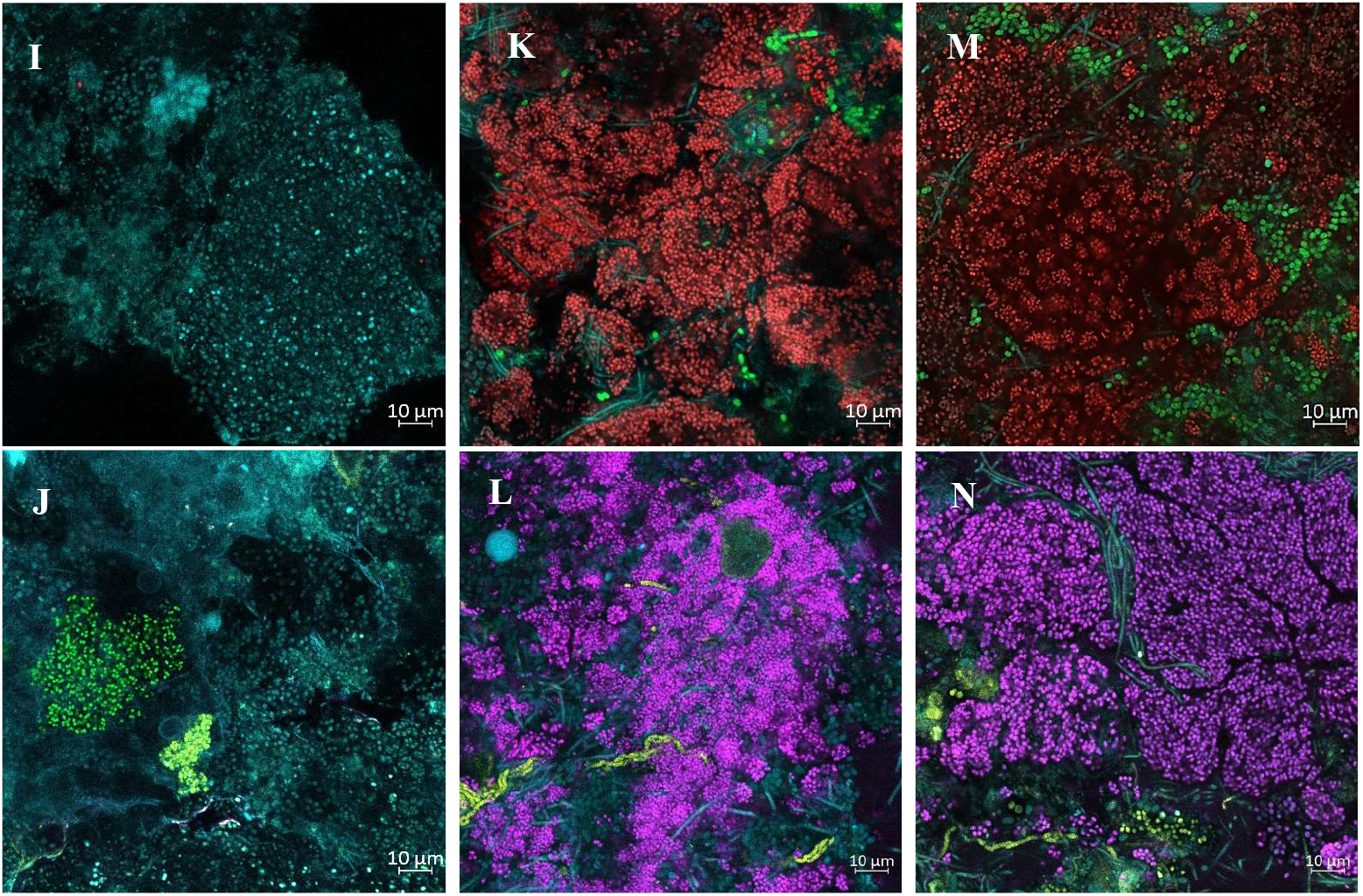
FISH images of key microorganisms in the EBPR system. (**A**) Seed sludge, (**C**) R30 on Day55, (**E**) R35 on Day55, (**G**) R30 on Day 119, (**I**) R35 on Day119, (**K**) R30 Day 294, (**M**) R35 Day 294, with EUBmix-Alex405 (EUB338, EUB338II and EUB338III) targeting all bacteria (Cyan), PAOmix-Cy3 (PAO651, PAO462 and PAO846) targeting *Ca.* Accumulibacter (Red), GBmix-FAM (GAO431 and GAO989) targeting *Ca.* Competibacter (Green). (**B**) Seed sludge, (**D**) R30 on Day55, (**F**) R35 on Day55, (**H**) R30 on Day 119, (**J**) R35 on Day119, (**L**) R30 Day 294, (**N**) R35 Day 294, with EUBmix-Alex405 targeting all bacteria (Cyan), PAOmix-Cy5 (PAO651, PAO462 and PAO846) targeting Ca. Accumulibacter (Purple), TFOmix-FAM (DF218 and DF618) (Yellow) targeting Cluster I *Defluviicoccus,* and DFmix-Cy3 (DF988, DF1020) targeting Cluster 2 *Defluviicoccus* (Green).

#### 3.2.3 PAO-GAO interaction regulation via controlled organic carbon supply benefited a higher EBPR stability at elevated temperature

For these core microorganisms in the EBPR community, we observed increases in relative abundance values of almost all ASVs in each lineage (both PAOs and GAOs) (Supplementary material, Fig. S4), and the development of originally undetectable rare species in the seed sludge in both reactors at the beginning of the operation, suggesting that the operation conditions employed in this work effectively benefited the growth of each community. These results also imply that the elevated temperature of 35°C is not an obstacle for a single ASV to survive and multiply. Compared to R30, the GAO community (*Ca.* Competibacter- and *Defluviicoccus-lineage,* and *Dechloromonas* ASVs 10 and 21) was relatively more and *Ca.* Accumulibacter was relatively less abundant in R35, which is in line with the lower EBPR performance observed. Within each lineage of GAOs and PAOs, significant dominance of respective ASVs was observed in R35 compared to R30, with lower community diversity (reflected by the Hill numbers ^1^*D*) and evenness (reflected by ^2^*D*, Supplementary material, Fig. S5), implying that both PAO and GAO communities were under stress at 35°C. Previous research suggested that both PAOs and GAOs have faster metabolic rates of carbon usage and increased maintenance coefficients when temperature increased (Brdjanovic et al., 1997; Lopez-Vazquez et al., 2007; 2008), which would result in increased carbon requirements of the EBPR community at 35°C compared to 30°C. We posited that the stress faced by the community at 35°C was largely caused by a deficiency in carbon sources. To test this hypothesis, the TOC/P ratio in R35 was increased from 25:1 to 40:1 on day 175 (Fig. 1). A subsequent increase in EBPR activity was observed together with increases in the diversity and evenness of all respective lineage members to values comparable to these in R30 (Supplementary material, Fig. S5), supporting the notion that supply of carbon was a limiting factor. The lower relative abundance of PAOs and higher abundance of GAOs at 35°C suggests that PAOs are more sensitive to carbon deficient conditions (low C/P ratios in the influent). Increasing the concentration of carbon sources effectively triggered *Ca.* Accumulibacter to outcompete *Ca.* Competibacter in terms of relative abundance by the end of the experiment (Fig. 2B and 2D).

There was no clear anticorrelation between relative abundance values of PAOs and GAOs in both reactors (Supplementary material, Fig. S6), implying that the variation in EBPR activities was not directly triggered by preferential growth of GAOs. Microbial ecological network analysis did not identify any direct interactions between PAOs and GAOs either (Supplementary material, Fig. S7). Previous research suggested that the superior (anaerobic) carbon uptake rates of GAOs provide an advantage at temperatures above 25°C. However, these apparent anaerobic carbon uptake rates were obtained under fast-feeding conditions, where carbon sources were fed into the reactor as a pulse (Lopez-Vazquez et al., 2007). To date, almost all laboratory-scale reactor experiments reported in the literature employed a fast-feeding regime. A modeling study also used the maximal anaerobic carbon uptake rates to evaluate PAO-GAO competition (Lopez-Vazquez et al., 2009). In full-scale WWTPs (Law et al., 2016; Qiu et al., 2019) and in this work the anaerobic carbon uptake was not determined by the maximal uptake rates of both organisms but by the feeding rate. The average carbon feeding rate was 3.21 mmol C/h/l in R30 and up to 5.15 mmol C/h/l in R35 (around 0.04 - 0.064 mmol C/mmol C MLSS), which is far below the maximum carbon uptake rates of 0.20 mmol C/mmol C for *Ca.* Accumulibacter and 0.30 mmol C/mmol C for *Ca.* Competibacter at 35°C (Lopez-Vazquez et al., 2007). Their actual carbon uptake rate was largely limited by the availability of carbon sources. The capability of *Ca.* Accumulibacter to generate luxury PMF via the efflux of protons in symport with PO_4_^3-^-P (Burow et al., 2008, Qiu et al., 2020) likely confers an advantage in competing with GAOs at low substrate concentrations.

### 3.3 PAO kinetics and stoichiometry are not compromised at elevated temperature

Previous research suggested that high temperature is unfavorable for EBPR mainly due to the lower anaerobic carbon uptake rates and the higher maintenance coefficient of *Ca.* Accumulibacter-PAOs compared to *Ca.* Competibacter-GAOs at temperatures above 20°C (Lopez-Vazquez et al., 2007). Regarding their aerobic metabolism, the aerobic carbon and P cycling kinetics of *Ca.* Accumulibacter continued to increase at the temperature range of 10-30 °C (Brdjanovic et al., 1997) while the aerobic kinetics of *Ca.* Competibacter showed no advantage over *Ca.* Accumulibacter (Lopez-Vazquez et al., 2008). On the contrary, the aerobic carbon conversion rates of *Ca.* Competibacter decreased significantly above 30°C with significantly reduced oxidative phosphorylation efficiency, reflected by the lower aerobic maximum yields, and higher PHA and oxygen requirements for glycogen production (Lopez-Vazquez et al., 2008). It is still not clear if the kinetics of aerobic metabolism in *Ca.* Accumulibacter continue to increase or if aerobic activity declines at temperatures above 30°C as it does for *Ca.* Competibacter. Additionally, the previous understanding was mainly obtained by exposing enrichment cultures obtained at 20°C to high temperatures. It is necessary to know if EBPR communities growing at higher temperatures have a similar response to temperature changes. To test previous observations and further understand the temperature effects on EBPR at temperature above 30, the P and carbon cycling activities in both reactors were monitored via consecutive cycle studies. In the first cycle the temperature chosen was that of the respective operational temperatures in reactors (30°C and 35°C for R30 and R35, respectively). In the next cycle, the temperatures of the two reactors were swapped (30°C for R35 and 35°C for R30). This design allowed an investigation of the temperature effects but avoided any bias due to the acute response of the community to temperature changes.

Consecutive cycle studies were performed five times throughout the long-term operation (on Days 55, 77, 105, 203 and 294) (Fig. 5 and Supplementary material, Fig. S8). Generally, higher P release was observed at 35°C in both reactors, since the amount of organic carbon input in the consecutive cycle was the same (Fig. 6). Increased P release suggested a higher maintenance requirement of PAOs (Lopez-Vazquez et al., 2007). But more pronouncedly, significantly higher P uptake and release rates were observed at 35°C than at 30°C in both reactors in all cases regardless of community composition, showing higher aerobic activities of *Ca.* Accumulibacter at 35°C, and this was not restricted to the presence of specific species/clades. Additionally, higher PHV formation was observed at 35°C than at 30°C in both reactors (Fig. 6). Different from *Ca.* Accumulibacter, glycogen served as both the reducing power source and the energy source for GAOs to drive acetate uptake (Filipe et al., 2001; Zeng et al., 2003; Oehmen et al., 2006). The reducing power generated during glycogenolysis and the pyruvate decarboxylation process is more than enough to reduce acetyl-CoA to hydroxybutyrate monomers. To balance the redox potential, around 1 in 4 parts of pyruvate is routed to the reductive branch of the TCA cycle to generate propionyl-CoA via the succinate-propionate pathway (Filipe et al., 2001; Zeng et al., 2003; McIlroy et al., 2014; Nobu et al., 2014), leading to the formation of PHV (around 24% of total PHA). The reduction of fumarate to succinate via the activity of fumarate reductase was also predicted to contribute to the generation of PMF for acetate uptake (Saunders et al., 2007; McIlroy et al., 2014). The higher PHV formation at 35°C indicates increased anaerobic activities of GAOs. This hypothesis is supported by the higher amount of glycogenolysis (Fig. 6). A previous study suggested a higher anaerobic carbon uptake rate of *Ca.* Competibacter when the temperature increased (Lopez-Vazquez et al., 2007). In view of the slightly higher relative abundance of *Defluviicoccus* in R35 than in R30, *Defluviicoccus* probably also has a similar temperature dependency. The aerobic carbon conversion related kinetics and stoichiometry parameters showed less significant increase as compared to the these in the aerobic P uptake related parameters, probably indicating the relatively lower aerobic carbon metabolism of GAOs than of *Ca.* Accumulibacter. This result is in line with the previous finding that the aerobic metabolic efficiency of *Ca.* Competibacter decreased at temperatures above 30°C (Lopez-Vazquez et al., 2008).

**Figure 5.**
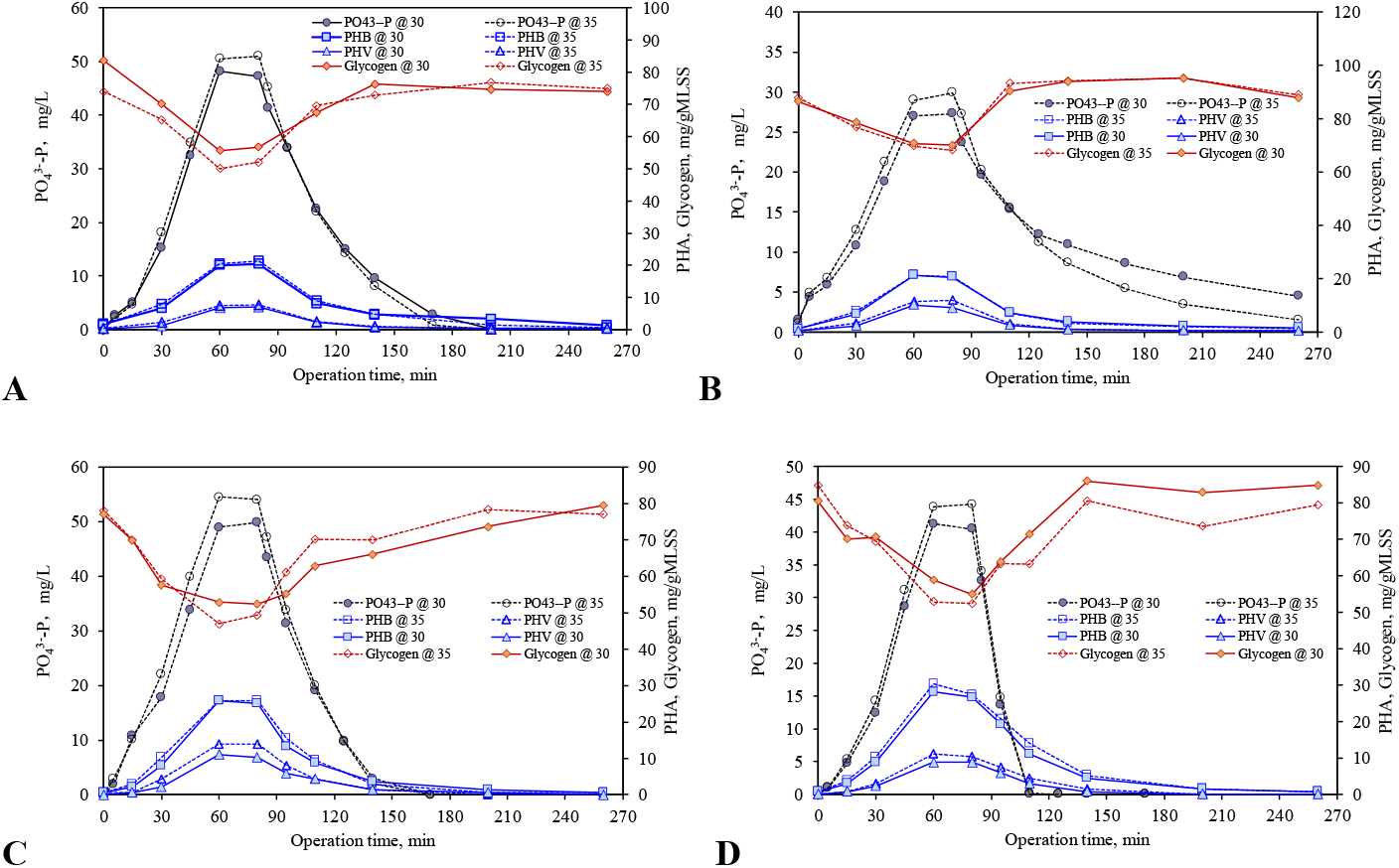
Representative carbon and phosphorus cycling characteristics in (**A**) R30 at Day77 and (**B**)R35 at Day77; and (**C**) R30 at Day 294 and (**D**) R35 at Day 294.

**Figure 6.**
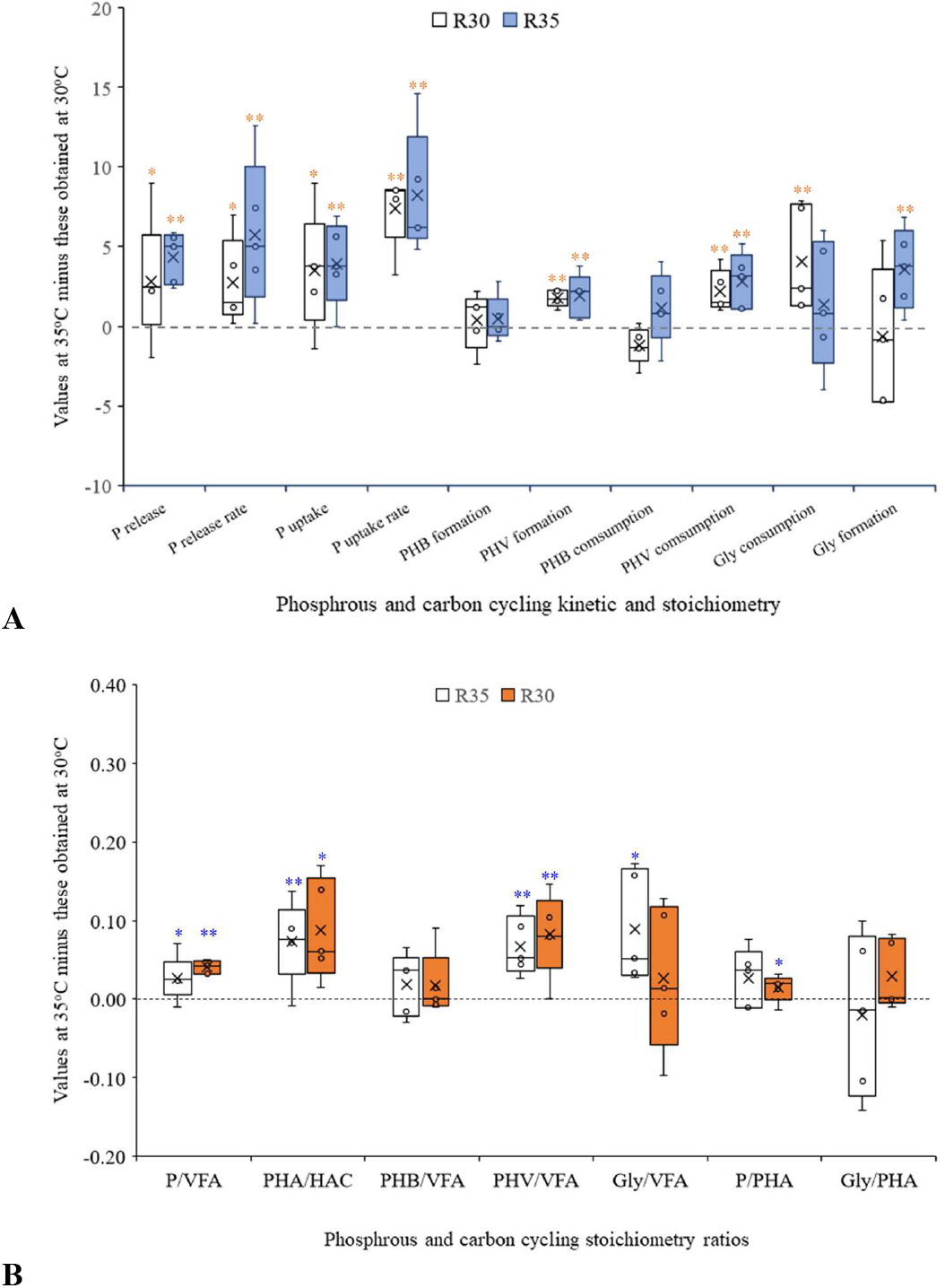
Differences in (**A**) the P and carbon cycling kinetic and stoichiometry parameters and (**B**) the stoichiometry ratios in the consecutive cycle study between 30°C and 35°C (n=5). Z-test was used to test if the mean difference values is significant different from 0 (* indicates P<0.05; ** indicates P<0.01).

In general, these results suggest that a temperature increase from 30°C to 35°C did not have a detrimental effect on the activity of PAOs. In contrast, the aerobic activity, especially P metabolism, increased significantly (Fig. 5 and 6). This advantage of *Ca.* Accumulibacter in aerobic metabolism may partially offset its weakness in anaerobic metabolism as compared to GAOs. Yet the increased anaerobic maintenance coefficient and the higher aerobic carbon consumption might have resulted in a carbon imbalance in the long-term, again underlining the importance of sufficient carbon supply to sustain long-term activity. These results explain the observation that *Ca.* Accumulibacter effectively co-existed with GAOs at 35°C and that a higher TOC/P ratio benefited its competitiveness vis-à-vis GAOs.

In addition to the consecutive cycle study, a normal cycle study was performed to monitor the changes in P and carbon cycling kinetics and stoichiometry during the long-term operation (Table 1). Canonical-correlation analysis (CCA) was used to relate these parameters to the community dynamics in both reactors (Fig. 7). Generally, the *Ca.* Accumulibacter population was positively related to the anaerobic P release/VFA uptake (P/VFA) ratios, the aerobic P uptake, the aerobic P uptake rate, and the aerobic P uptake/PHA consumption (P/PHA) rates; these parameters are effective indices of a higher EBPR activity. The population of GAOs was positively related to the anaerobic VFA uptake/PHA formation (VFA/PHA) and glycogen consumption/VFA uptake ratios, higher values of which are indications of the higher relative abundance values of GAOs. *Dechloromonas* was also positively related to these factors, implying that they probably performed as GAOs. In contrast to the high correlation between the *Ca.* Accumulibacter population and the P uptake rate, a low correlation was observed between the *Ca.* Accumulibacter population and the P release rate. In both systems, a slow-feeding strategy was used, limiting the P release and C uptake rates of *Ca.* Accumulibacter. The strategy also effectively limited the advantage of *Ca.* Competibacter in anaerobic C uptake at high temperature. We estimated the percentage of C taken up by PAOs in both systems based on the anaerobic P/VFA ratios and aerobic P/PHA ratios. Our previous research suggested that the P/VFA ratio of a highly enriched *Ca.* Accumulibacter culture at 30°C is in the range of 0.51 - 0.59 (Qiu et al., 2020). And based on the metabolic model, the P/PHA ratio of *Ca.* Accumulibacter is around 0.42. By comparing the anaerobic P/VFA and aerobic P/PHA ratios observed in this work and the model values (Table 1) around 70% (59.6 - 91.9% with an average of 76.3% based on the P/VFA ratios, or 44.3 - 79.7% with an average of 64.7% based on the P/PHA ratio) of the C sources was taken up by *Ca.* Accmulibacter in R30. In R35 it was around 50% (33.8 - 73.1% with an average of 56.8% based on the P/VFA ratios, or 27.9 - 66.5% with an average of 44.0% based on the P/PHA ratio) with the highest values observed at high TOC/P ratios (Fig. 8), showing that the slow-feeding strategy could effectively prevent excessive uptake of C sources by GAOs even at high TOC/P ratios.

**Table 1.**
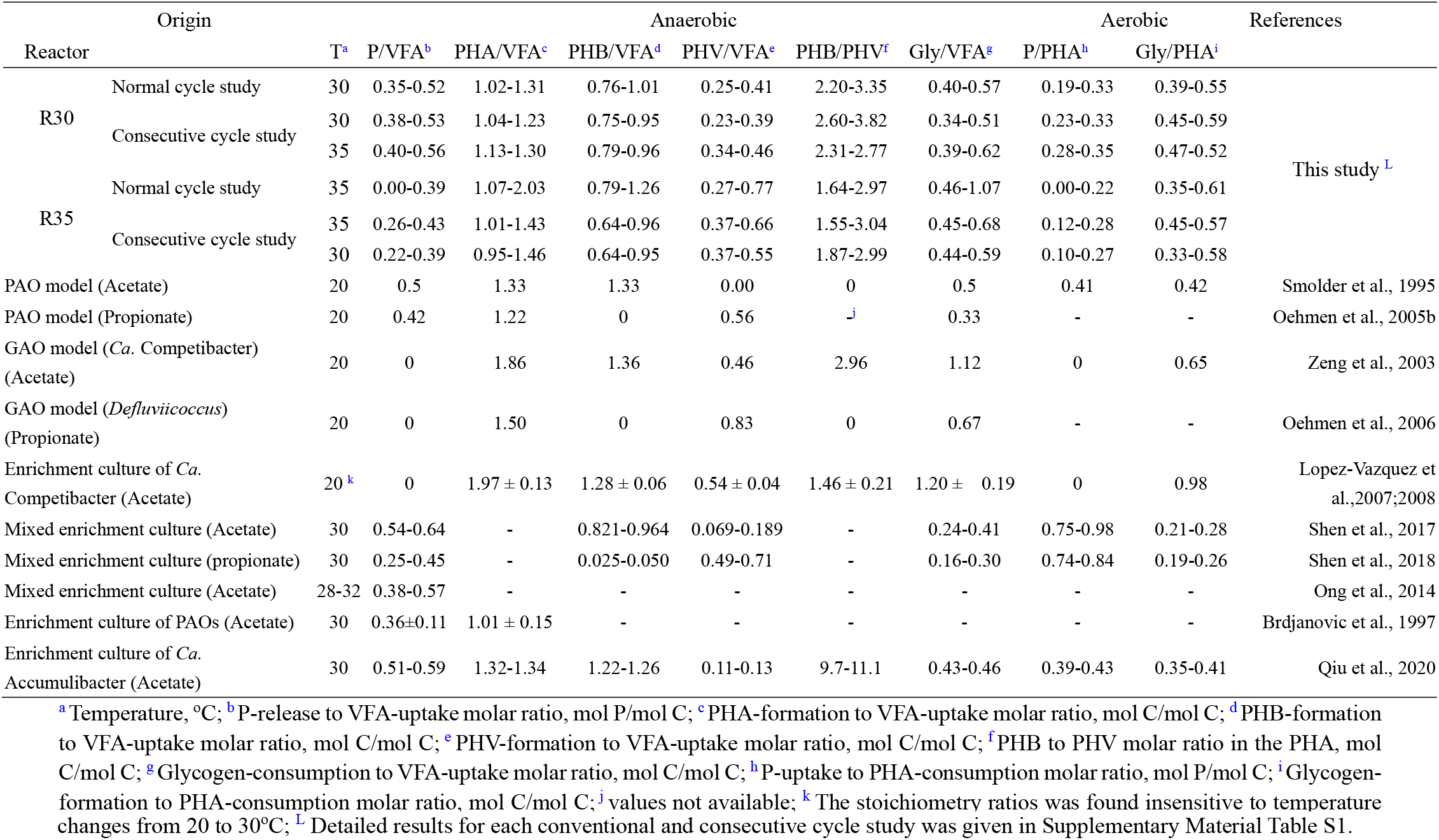
P and carbon transformation stoichiometry ratios obtained in this study and a comparison with literature and model values

**Figure 7.**
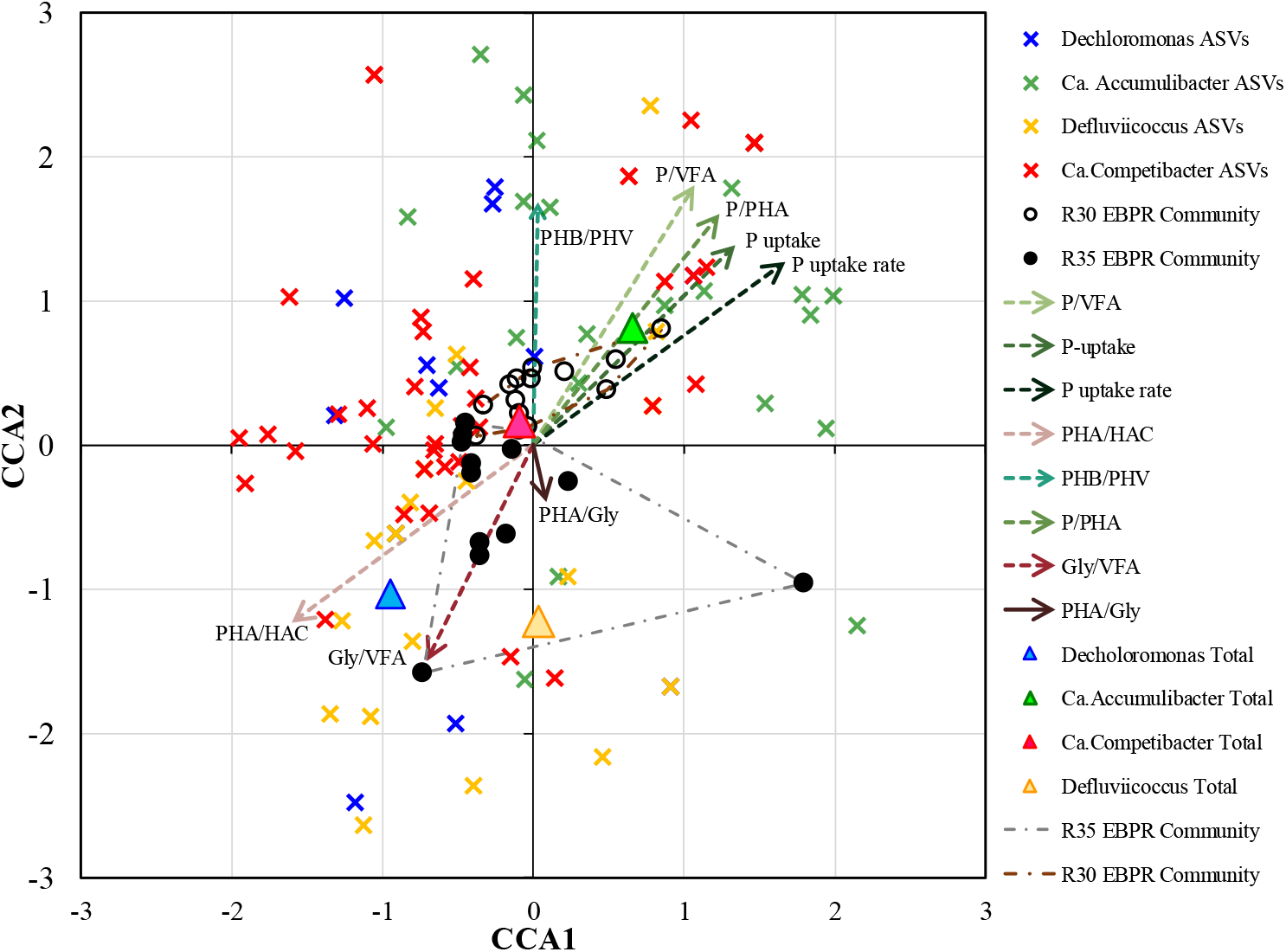
CCA analysis revealing the relationship between the EBPR community and the kinetic and stoichiometry parameters.

**Figure 8.**
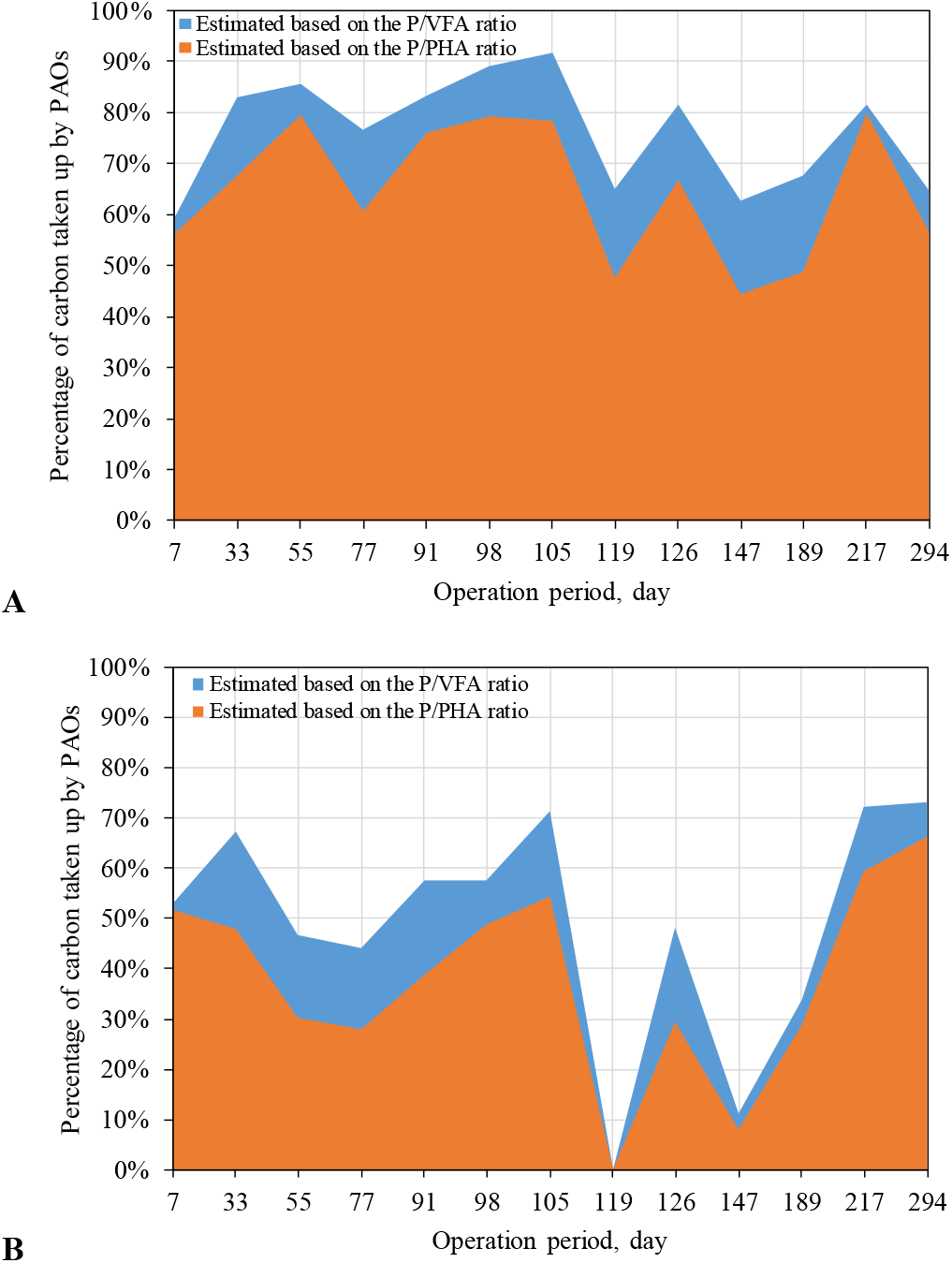
Estimated percentage of carbon taken up by PAOs in (**A**) R30 and (**B**) R35.

## 4 CONCLUSIONS

The following findings will have important implications for a successful operation of EBPR systems at elevated temperatures:

- EBPR at 35°C is feasible. A wide range of *Ca.* Accumulibacter ASVs closely related to those commonly found in full-scale and lab-scale EBPR systems operated at temperate conditions could survive and proliferate for a prolonged period (over 300 days), suggesting that EBPR at elevated temperature does not require a highly specialized EBPR community.
- Low biodiversity and evenness were observed at 35°C for each specific PAO/GAO linage, showing that the EBPR community was stressed as a consequence of higher C (and/or P) metabolic rates and a resultant carbon deficiency. Increasing the TOC/P ratio, eased the community competition and benefited EBPR performance at 35°C. Increased carbon input might be necessary for stable EBPR at this temperature.
- Short-term temperature tests showed increased activity of *Ca.* Accumulibacter (especially increased P uptake rates) when temperature increased from 30°C to 35°C, suggesting that elevated temperature does not have a direct adverse effect on *Ca.* Accumulibacter.
- Slow-feeding strategy effectively limits the carbon uptake rates of GAOs, thus allowing *Ca.* Accumulibacter to co-exist with GAOs and outcompete them at 35°C to achieve complete P removal.
- Cluster III *Defluviicoccus* members could effectively survive the slow-feeding condition, implying that their bioenergetic characteristics for C uptake are different from those of their tetrad-forming relatives.

## Acknowledgements

This research was supported by the Singapore National Research Foundation and the Ministry of Education under the Research Centre of Excellence Programme, and by a research grant from the National Research Foundation under its Environment and Water Industry Programme (project number 1102–IRIS–10–02), administered by PUB-Singapore’s national water agency. Dr. Guanglei Qiu acknowledges the support of National Natural Science Foundation of China (No. 51808297).

## Notes

### Competing Interest Statement

The authors have declared no competing interest.

