## Supplementary Materials for "Global warming readiness: Feasibility of enhanced biological phosphorus removal from wastewater at 35°C"

**A**
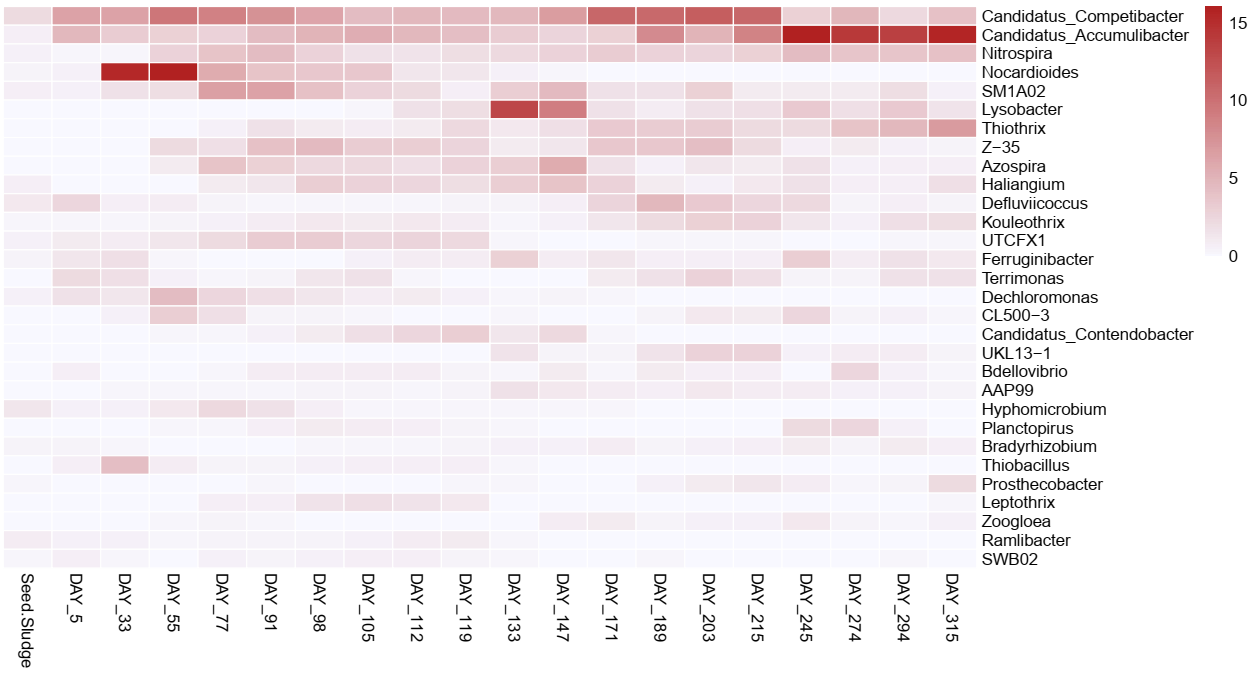

**B**
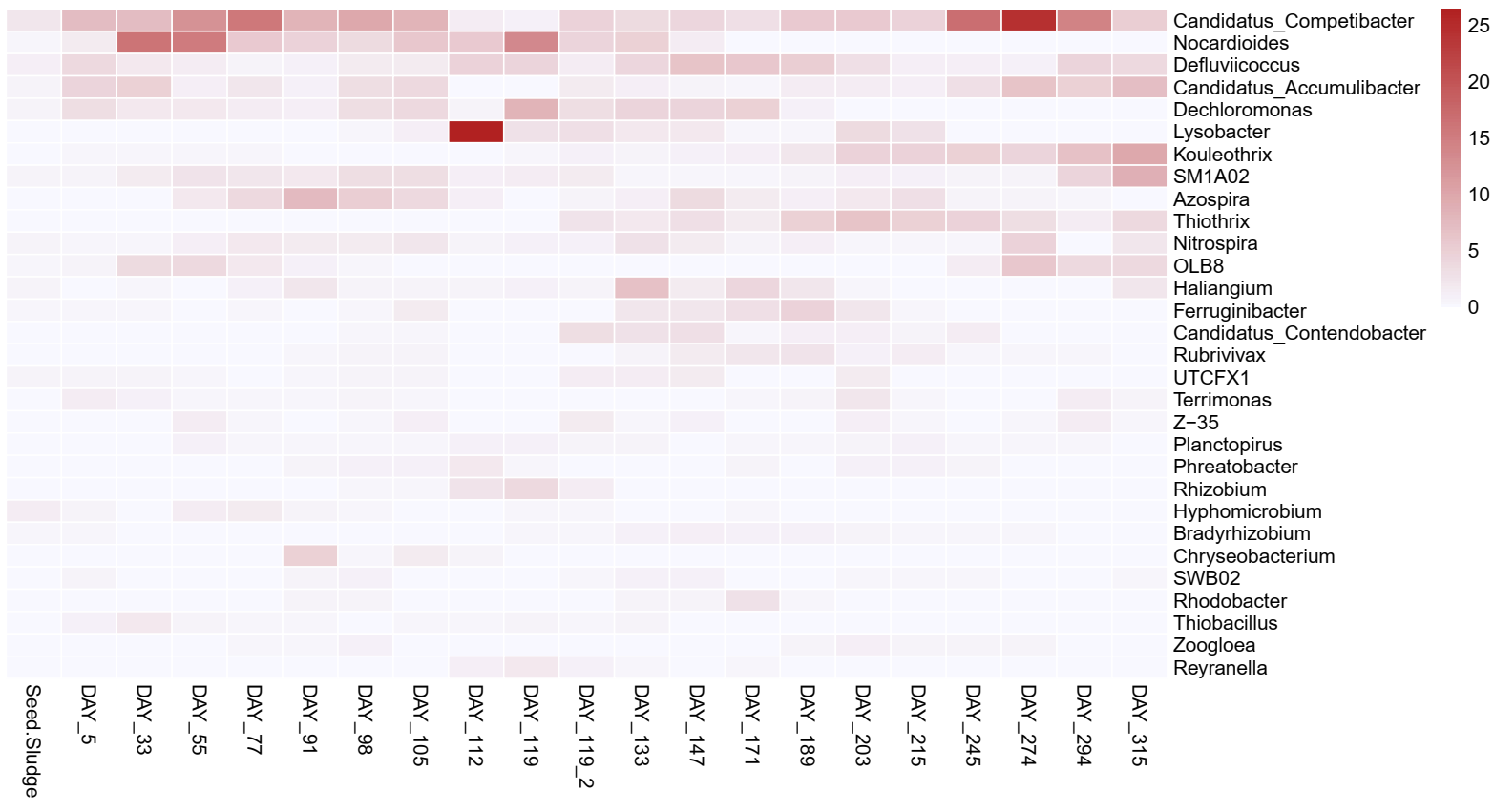

**Figure S1.** Top 30 most abundant genus and their dynamics along the operation of (**A**) R30 and (**B**)R35.

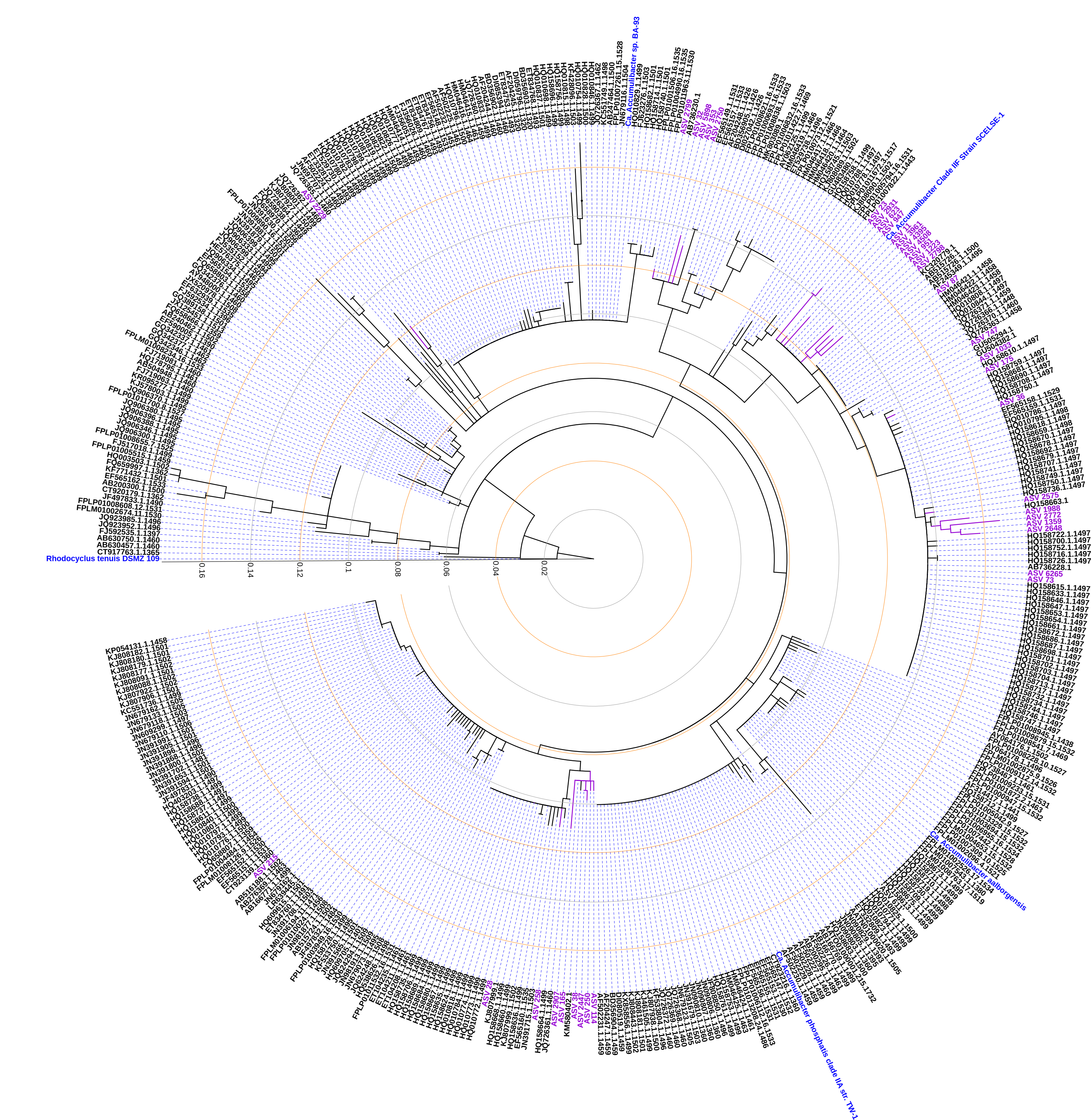

**Figure S2.** Phylogenetic analysis of the ASVs retrieved in the 16S rRNA amplicon analysis. *Rhodocyclus tenuis* DSMZ109 was used as an outgroup. The tree was generated with MEGA 10 using the maximum likelihood method. The tree was visualized using an online tool: iTOL (https://itol.embl.de).

A
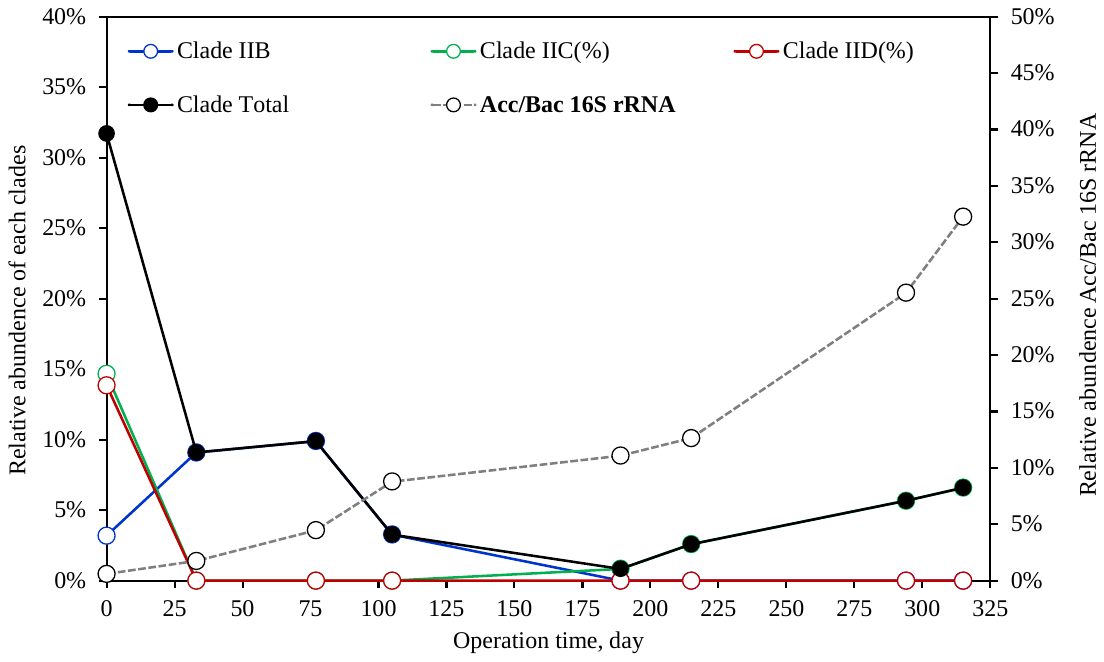

B
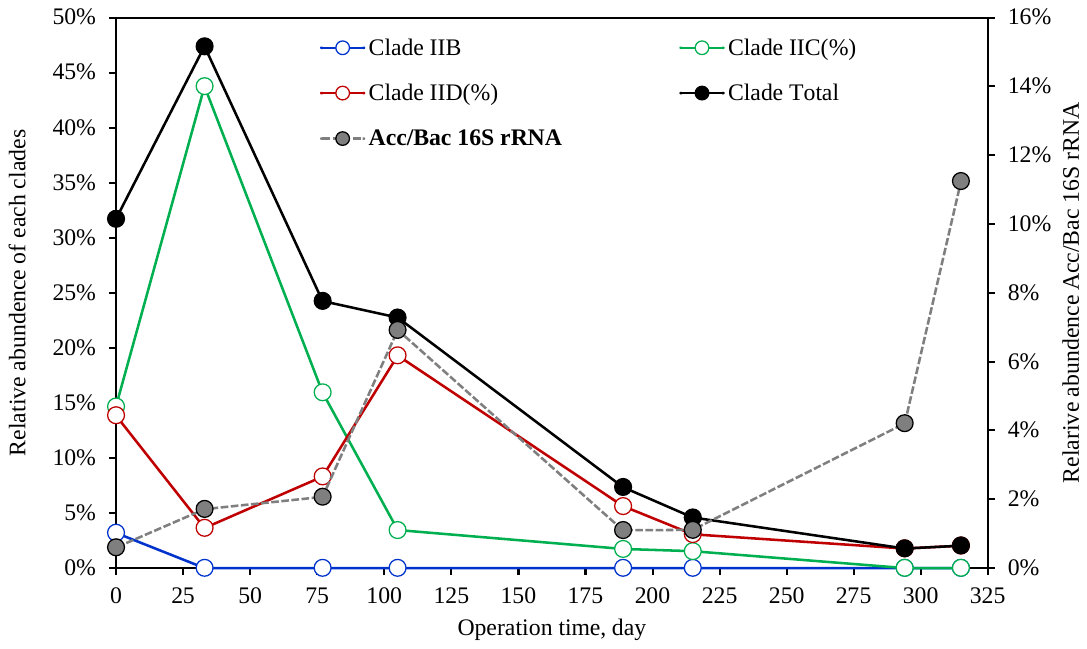

**Figure S3.** Clade level composition and dynamics of Ca. Accumulibacter in (**A**) R30 and (**B**) R35.

**A**
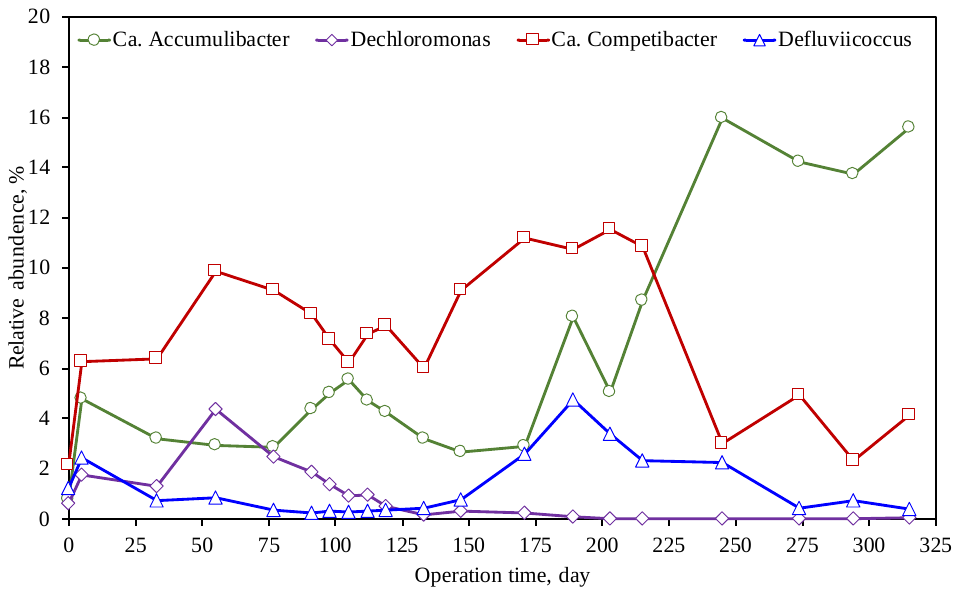

**B**
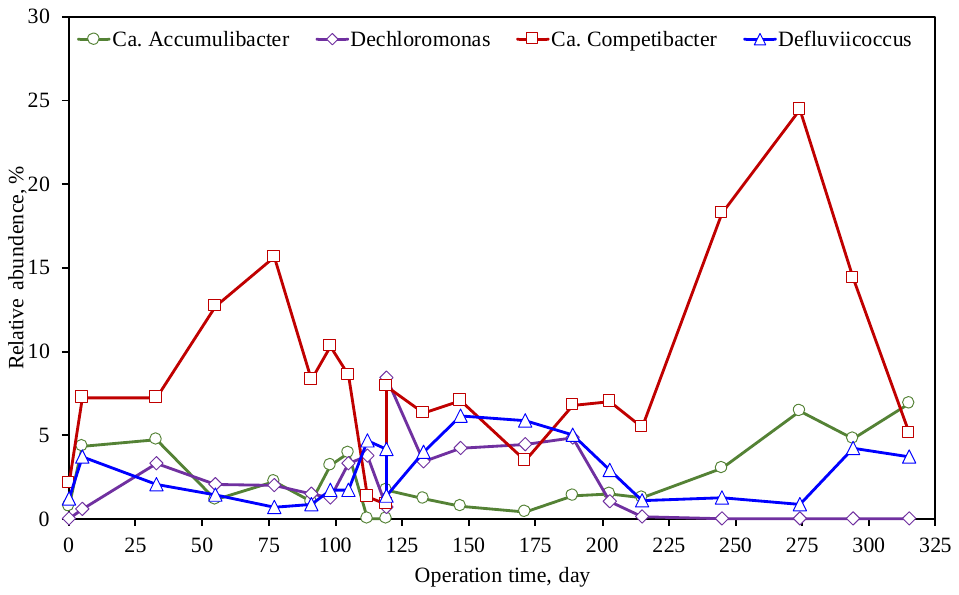

**Figure S4.** Dynamics in the key EBPR-related community in (**A**) R30 and (**B**) R35 revealed by 16S rRNA gene amplicon sequencing.

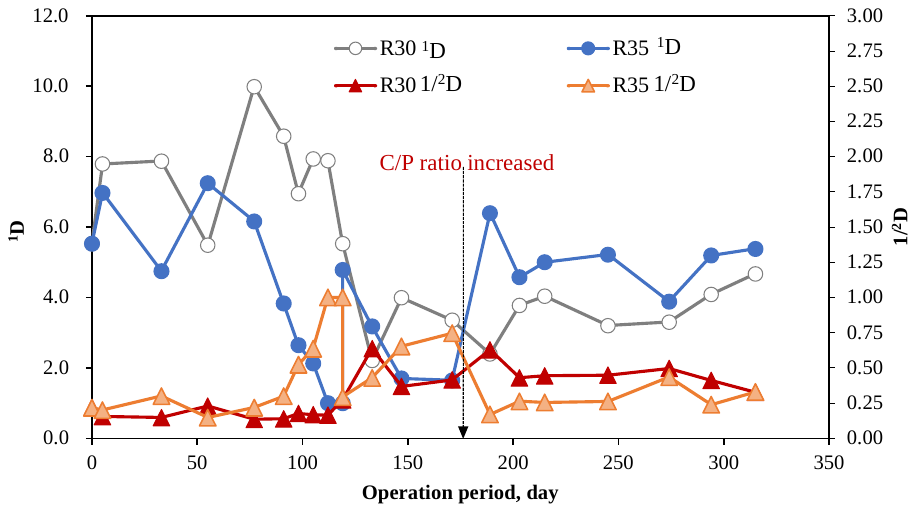

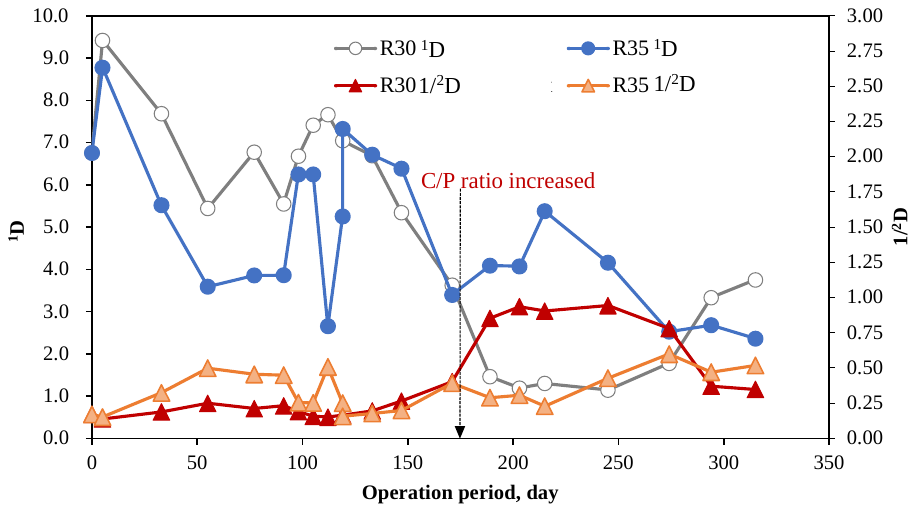

**Figure S5.** Changes in the Alpha diversity (Hill numbers: *^1^D* and *^2^D*) of the (**A**) *Ca.* Accumulibacter and (**B**) *Ca.* Competibacter communities in R30 and R35.

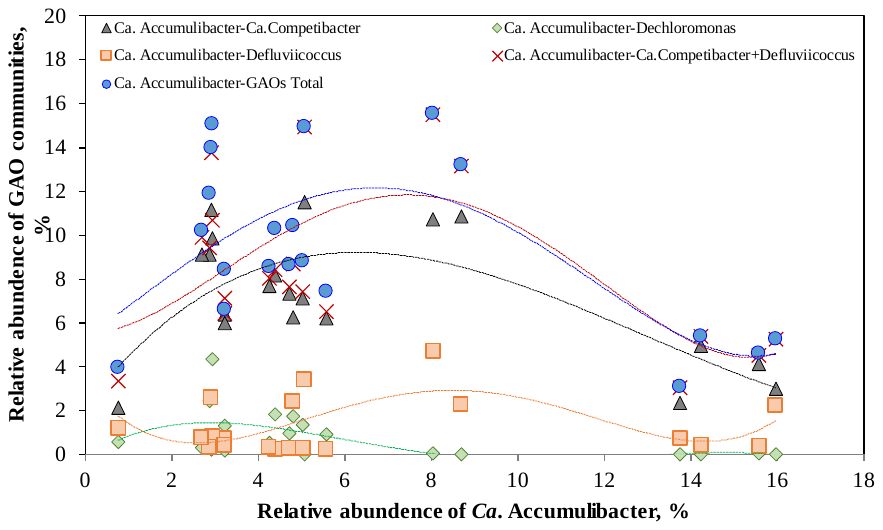

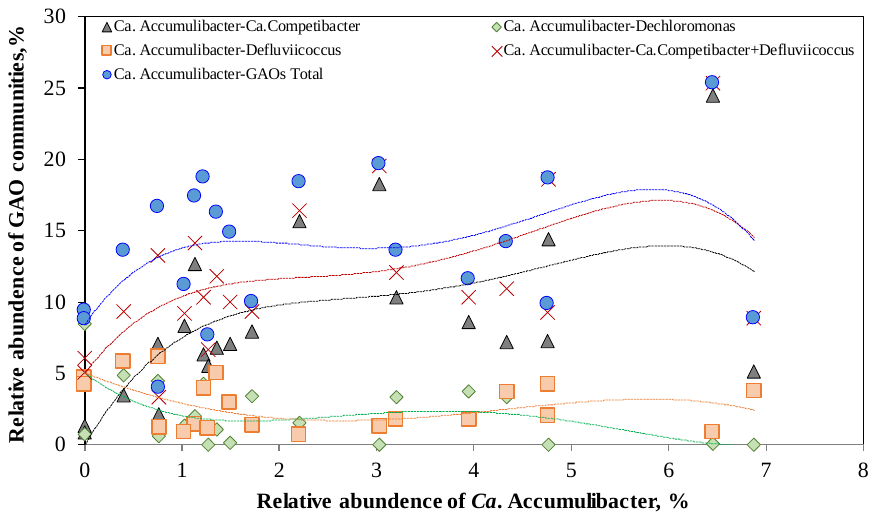

**Figure S6.** Correlation among the relative abundances of *Ca.* Accumulibacter and GAOs (*Ca.* Competibacter, Defluviicoccus, and Dechloromonas) communities in (**A**) R30 and (**B**) R35.

**A**
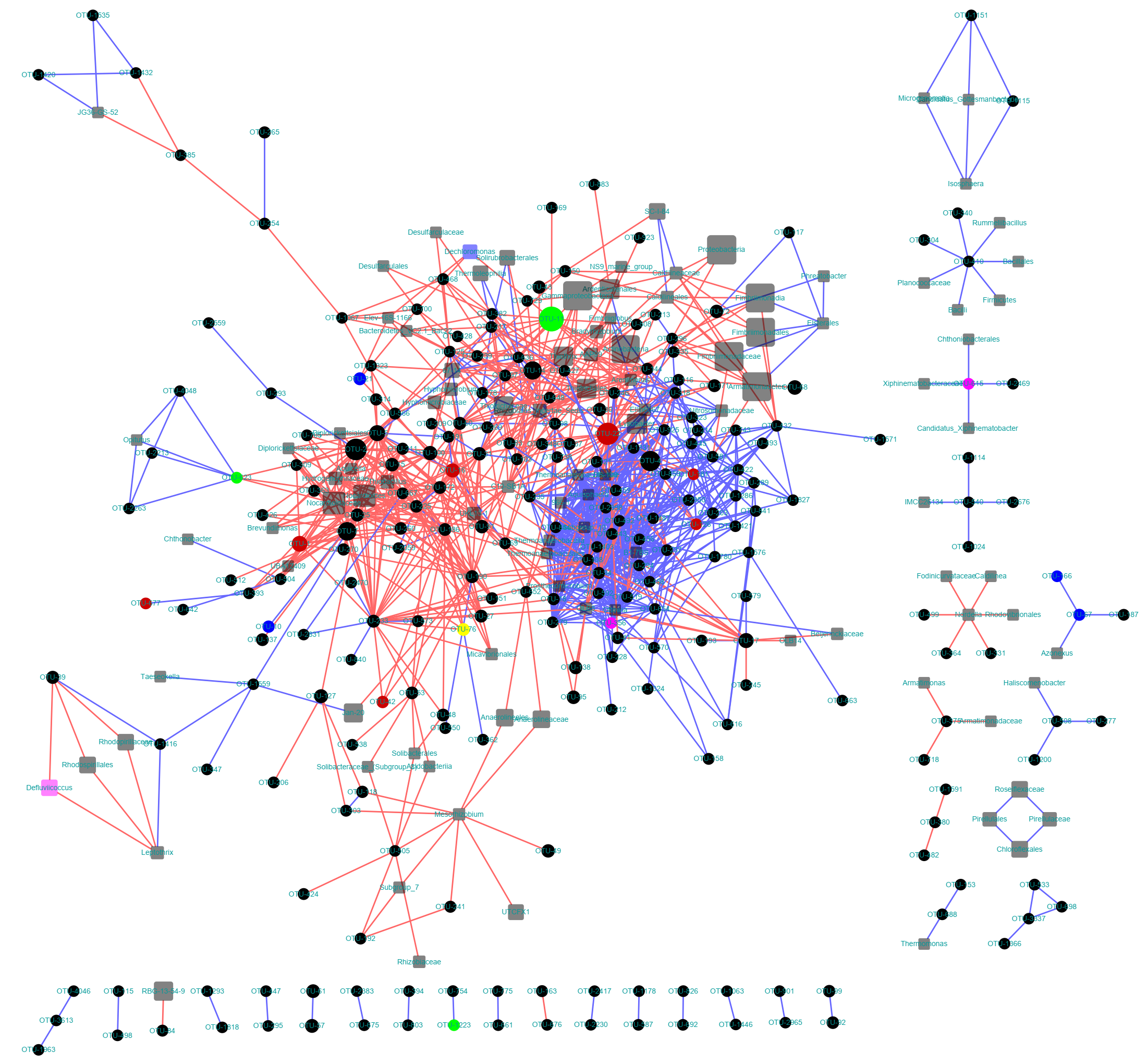

**B**
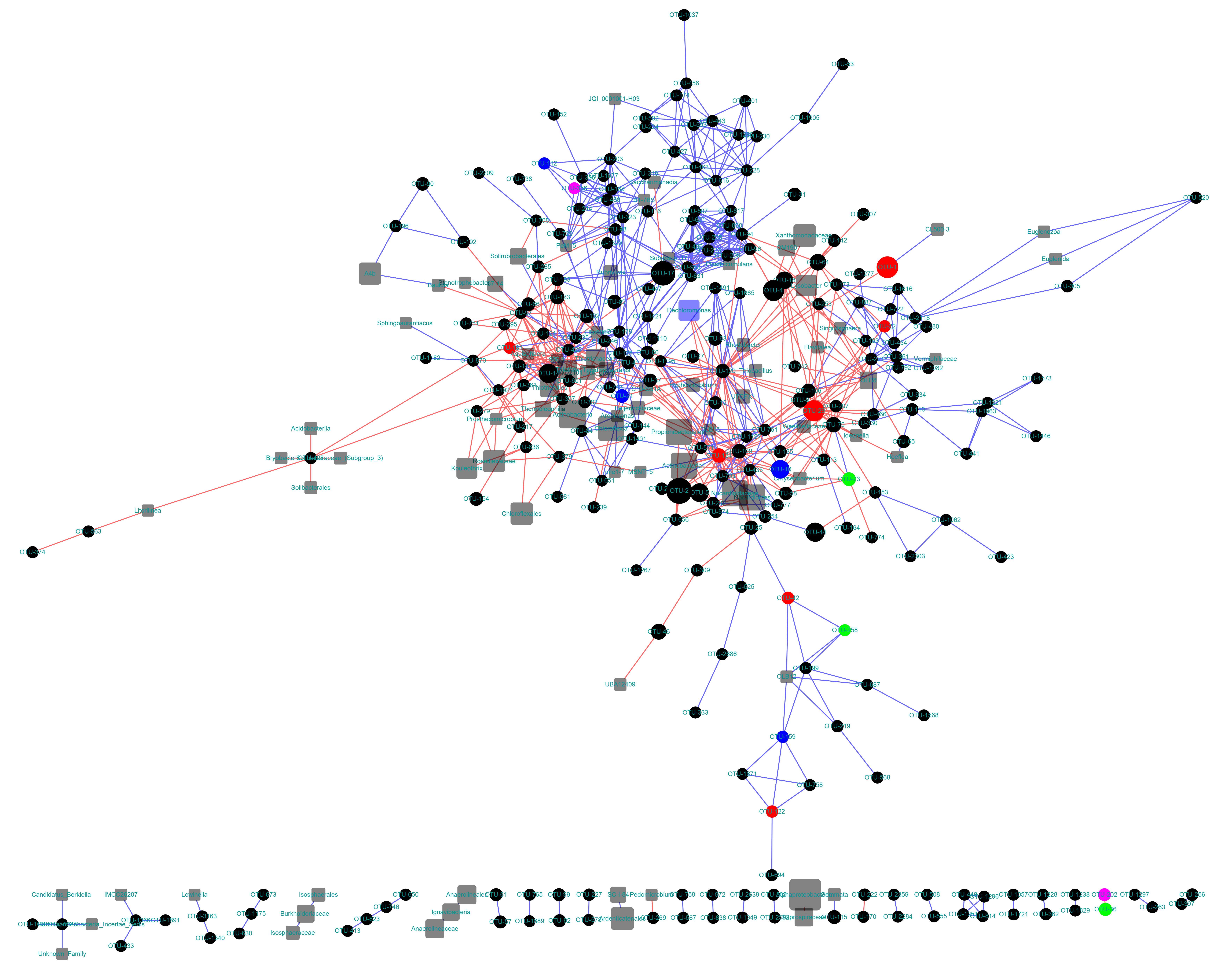

**Figure S7.** Microbial ecological network analysis showing the interactions between community members during the operation of the reactors: (**A**) R30 and (**B**) R35. The size of the solid cycle nodes indicated the relative abundance of the ASVs. Ca. Accumulibacter-, Ca. Competibacter-, *Defluviicoccus-*, and *Dechloromonas-* related ASVs are denoted in green, red, purple, and blue, respectively. The higher-level taxa are denoted as gray squares.

**A**
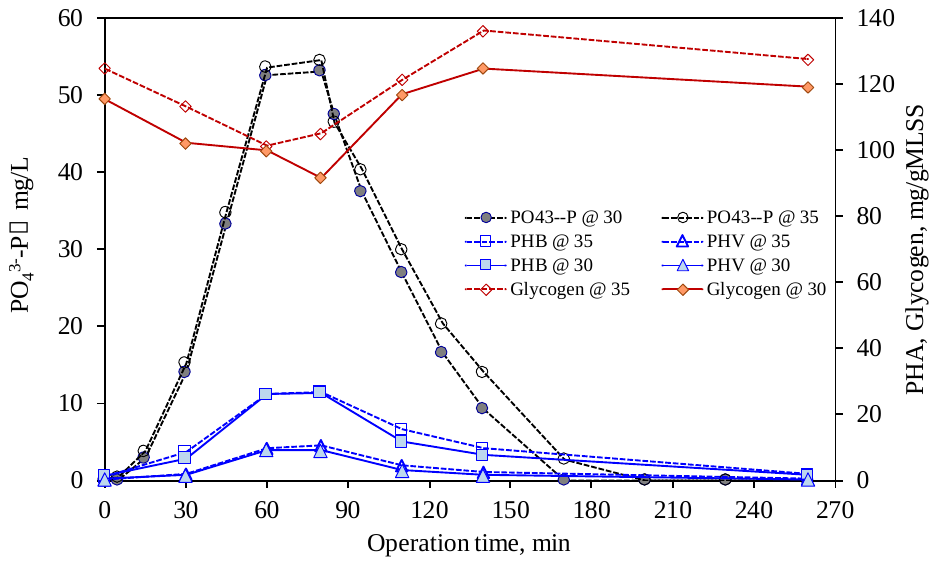
**B**
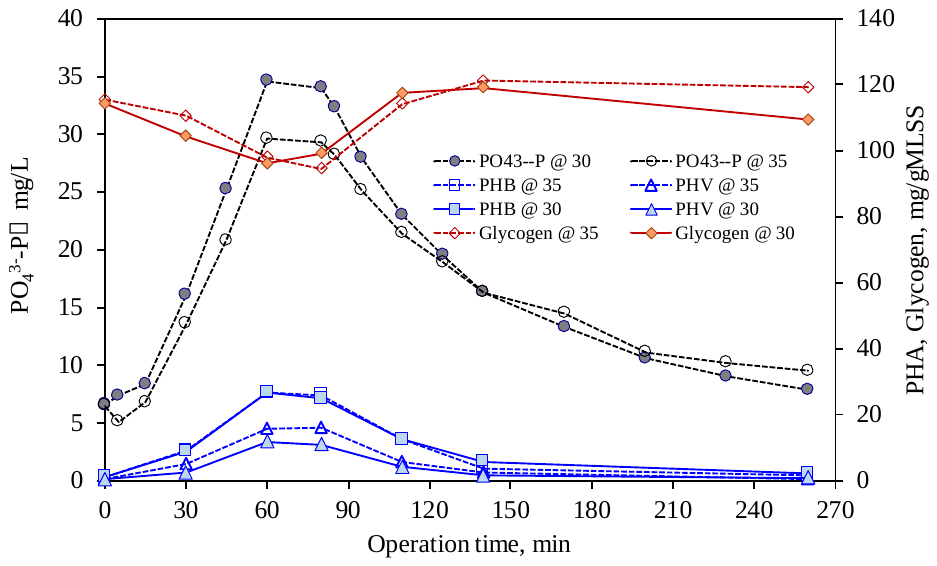

**C**
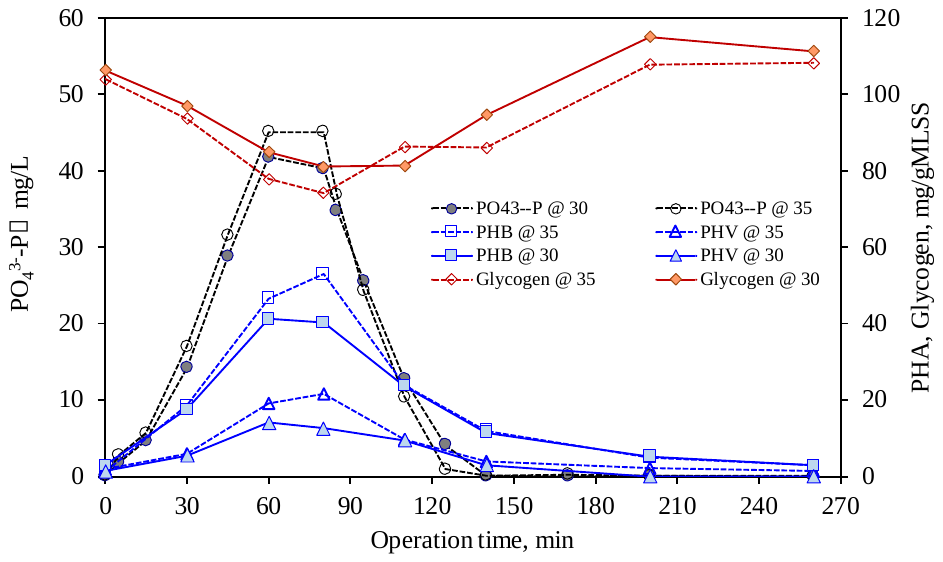
**D**
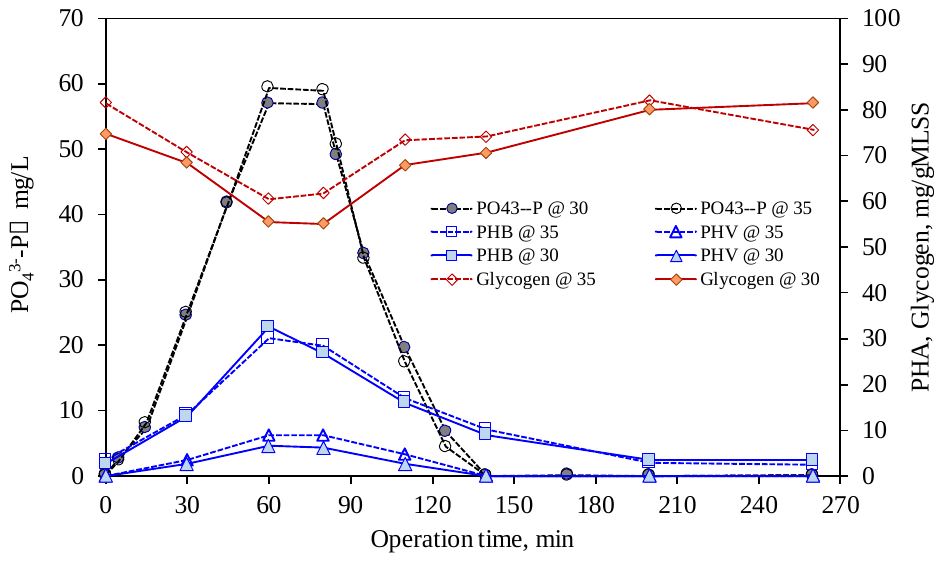

**E**
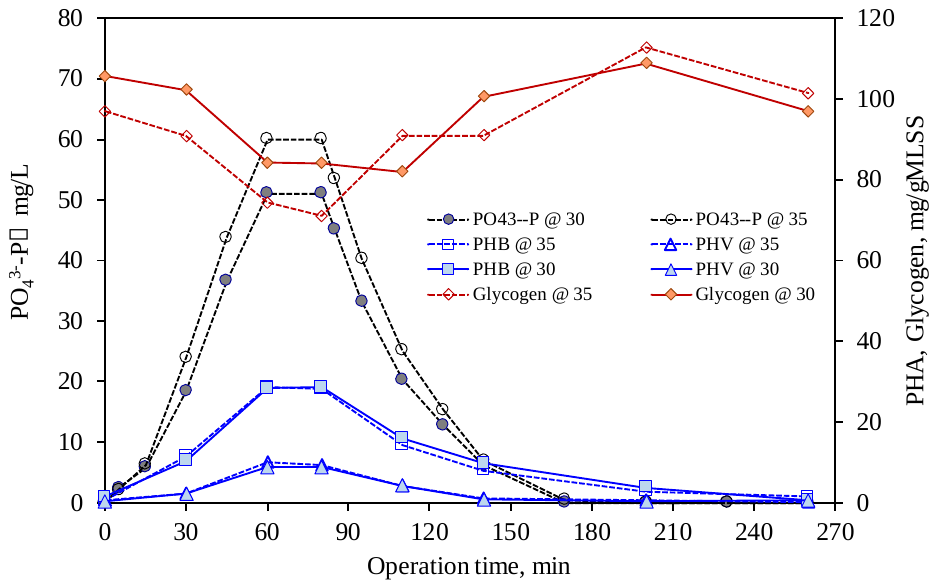
**F**
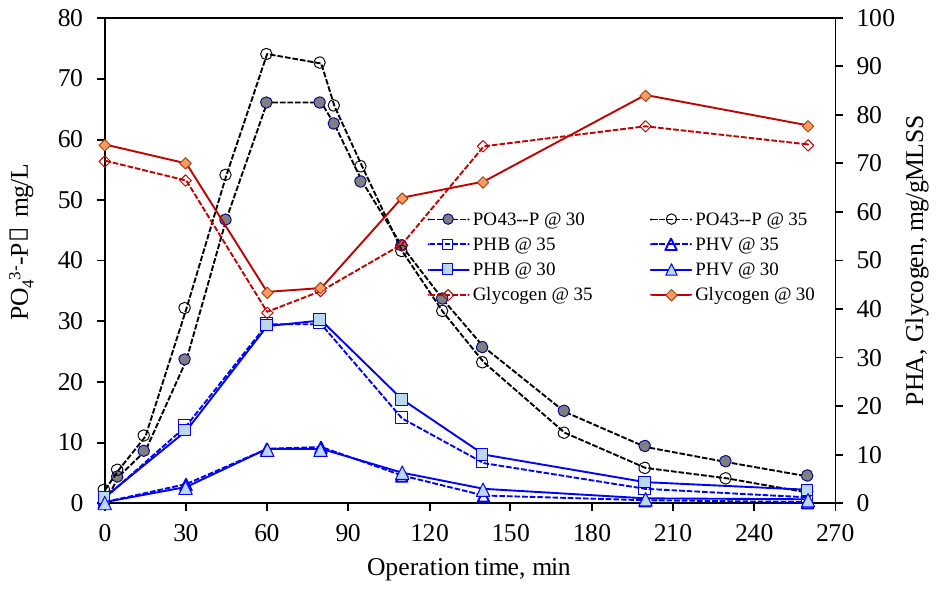

**Figure S8.** Representative carbon and phosphorus cycling characteristics in (**A**) R30 at Day55 and (**B**)R35 at Day55; (**C**) R30 at Day105 and (**D**) R35 at Day105; and (**E**) R30 at Day215 and (**F**) R35 at Day215.
